# OTU–Taxa: a taxonomy-aware pretrained model for 16S microbiome profiles

**DOI:** 10.64898/2026.11.23.768542

**Authors:** Hernan Perez, Ricardo Henao, Robert Hoehndorf

## Abstract

**Background:** Microbiome profiles combine sequence-derived units with taxonomy. Pretrained models use different community vocabularies and are often evaluated on few labels, leaving their value under broad competitive evaluation unclear. We introduce OTU–Taxa, which pairs operational taxonomic unit (OTU) identity with hierarchical taxonomy during autoregressive pretraining on unlabeled profiles.

**Results:** In an in-domain evaluation across 250 Microbe Atlas Project (MAP)-derived classification tasks with disjoint folds, mean area under the receiver operating characteristic curve (AUROC) was 0.744 for OTU–Taxa and 0.749 for Random Forest. OTU–Taxa was the highest-ranked pretrained method; Nemenyi comparisons did not detect rank differences from Random Forest across the three metrics. The model also generated largely non-copied profiles and recovered more withheld community members at larger cutoffs, while retrieval gave stronger empirical fidelity and early ranking. It also supported abundance-rank readouts and recovered higher-rank classifier assignments from sample context for known-vocabulary OTUs under controlled sequence degradation, although agreement was limited at genus and species ranks.

**Conclusions:** Within the MAP 97% closed-reference 16S vocabulary, the pretrained representation supports in-domain prediction of metadata labels alongside generative and taxonomic readouts. Supervised abundance models, empirical retrieval, and intact-sequence classifiers remain stronger choices for several corresponding endpoints, while OTU–Taxa provides one shared contextual representation across all four analyses.

## 1 Background

Microbial communities span host-associated and free-living environments and contribute to ecosystem and host states [1, 2]. In these settings, community members show non-random co-occurrence, exchange metabolites, and participate in trophic and syntrophic interactions that shape assembly [3–5]. In host-associated systems, shifts in community composition are also associated with disease phenotypes such as inflammatory bowel disease (IBD) [6, 7]. These community-level regularities motivate whole-profile models in which a member’s identity and abundance rank are interpreted relative to the observed community.

Shotgun metagenomics profiles community genetic content and functional potential, whereas 16S ribosomal RNA (rRNA) gene amplicon sequencing enables costeffective surveys across many samples [8, 9]. Public resources now support large-scale representation learning. The Microbe Atlas Project (MAP) harmonizes more than 1.8 million samples using common small-subunit (SSU) rRNA references and operational taxonomic units (OTUs) across studies and environments [10]. At this scale, the profiles are sparse, compositional, heterogeneous over studies and protocols, and tied to sequence-derived units whose taxonomic labels vary in depth [11–13]. In standard marker-gene workflows, feature-table rows correspond to units such as OTUs or amplicon sequence variants (ASVs), which can also be assigned taxonomy at broader ranks [13, 14]. The feature-level view preserves fine-grained observations of community composition, whereas taxonomic labels provide a structured vocabulary to relate features in broader biological ranks [13, 15, 16].

Large unlabeled biological datasets support reusable pretraining before task-specific adaptation in language, single-cell transcriptomes, and other biological profiles [17–20]. In microbiome data, recent pretrained models have explored OTU, taxonomic, metagenomic species, and phylogeny-aware vocabularies [21–24]. Pope’s model represents samples as OTU sequences, whereas the Microbial General Model (MGM) and Waypoint use abundance-ranked genus-level taxonomy profiles; Waypoint additionally falls back to higher-rank taxonomy tokens when genus-level assignments are unavailable [25–27]. In addition to abundance-ranked sequence formulations, recent work has explored permutation-invariant set representations that incorporate relative abundance during sequence-embedding aggregation [28]. Together, these approaches show that both the community vocabulary and the encoding of abundance and sample order are modeling choices, not just preprocessing details.

Although pretrained microbiome models have shown promising transfer in selected tasks and datasets [25, 26], their practical advantage is not yet settled under broader and more competitive evaluation. Recent benchmarks indicate that performance is strongly dependent on task, representation, and baseline considered, with classical supervised methods remaining strong in several microbiome prediction settings [22, 24, 27, 29]. This raises two connected questions: what reusable community signal is learned by pretraining on paired OTU identities and taxonomic annotations, and in which analyses is that signal competitive with or distinct from task-specific abundance models, empirical retrieval, or sequence-only taxonomy classifiers?

To study these questions, we introduce OTU–Taxa, a pretrained taxonomy-aware microbiome model that represents each sample as an abundance-ranked sequence of paired OTU identity and taxonomy tokens. Sample classification tests in-domain prediction of metadata labels across 639 MAP-derived tasks, of which the 250 tasks evaluated with disjoint folds form the primary benchmark; generative community readouts test learned community dependencies, and context-based taxonomy-assignment recovery under controlled sequence degradation tests whether sample context remains informative as local sequence evidence weakens. Together, these analyses test the paired model, identify tasks where empirical or supervised baselines remain stronger, and examine which outputs reflect community context.

## 2 Methods

After pretraining, we evaluate metadata-label prediction, generative readouts, and taxonomy inference from sample context.

### 2.1 Microbiome data sources and sample representation

We obtained amplicon sequencing profiles from MAP [10], which aggregates microbiome studies in a closed-reference OTU framework. We used the filtered MAP collection available as of August 2025, comprising 1,836,250 samples across diverse host and environmental contexts. Each sample is initially represented as a sparse vector of OTU identifiers with associated raw read counts.

MAP provides OTU definitions at multiple sequence-identity thresholds. We used the widely used 97% level as a single intermediate-resolution vocabulary, which yields 62,200 OTUs and balances biological resolution, vocabulary size, and sparsity across the heterogeneous corpus. Exact sequence variants provide finer resolution [13], whereas using an established clustering level provides a consistent corpus-wide representation without requiring separate model vocabularies for every available resolution.

We represent each microbiome sample as an abundance-ranked sequence of observed OTUs. OTUs represented by fewer than four reads in a sample are removed before ranking. The remaining OTUs are ordered by descending raw read count; OTUs with equal counts retain their order in the corresponding MAP source record. The ordered OTUs are converted to token identifiers from a global OTU vocabulary V_otu_. For a sample containing *L_x_* observed OTUs, we write the OTU sequence as **o***_x_* = (*o*_1_*, o*_2_*, . . . , o_Lx_* ), with *o_i_* ∈ V_otu_, where position *i* corresponds to the abundance rank within the sample. The general representation idea is inspired by rank-based biological-profile tokenization in models such as Geneformer [18]. Abundance is encoded only by token order; raw counts are not model inputs. Exact-tie handling and order-sensitivity controls are described in Additional file 1: Supplementary Methods M1A and M1E. Application to another dataset therefore requires its features to be mapped to the same 97% MAP closed-reference OTU identifiers. ASVs or OTUs that cannot be mapped to this vocabulary have no learned input token and cannot be used directly by the released checkpoint.

For Transformer batching, sequences are truncated or padded to a fixed maximum length *L* = 500. With one position reserved for the beginning-of-sample token, this cutoff retains the complete observed OTU set for approximately 95.5% of filtered MAP samples while controlling the quadratic memory cost of self-attention. Padding is excluded from losses and evaluation; *m_i_* is 1 for a non-padding position and 0 otherwise.

Each OTU token is paired with a taxonomy token. Let V_tax_ denote the taxonomynode vocabulary and let *ϕ* map an OTU to its assigned taxonomy node. For each non-padding OTU, *t_i_* = *ϕ*(*o_i_*), with *t_i_* ∈ V_tax_. A sample is therefore represented as aligned OTU–taxonomy pairs **x** = ((*o*_1_*, t*_1_), (*o*_2_*, t*_2_)*, . . . ,* (*o_Lx_ , t_Lx_* )). Unless otherwise stated, the pretrained models reported in the main experiments use this filtered MAP corpus without downstream sample labels; metadata labels are introduced only inside the downstream cross-validation folds.

The main model uses taxonomy labels assigned to each OTU representative sequence with the SINTAX classifier [30], as implemented in VSEARCH [31], using the SINTAX-formatted SILVA v123 16S reference distributed by USEARCH [15]. Taxonomic labels are assigned from kingdom through species. A rank is accepted only when the classifier’s confidence exceeds the default 80% threshold; otherwise, that rank and all lower ranks are treated as unresolved.

The taxonomy vocabulary is built from the resulting hierarchy. The mapping *ϕ* : V_otu_ → V_tax_ assigns each OTU to its deepest accepted taxonomy node. Each taxonomy token is therefore an annotation of its paired OTU, providing a hierarchy-structured view of the same sequence-derived feature. In the SINTAX-derived full MAP reference, this procedure produced 6,929 taxonomy nodes.

We selected SINTAX [30] because it is widely used for amplicon taxonomic assignment, does not require fitting a new classifier for the study-specific reference set, and scales to repeated model training and benchmarking.

### 2.2 Autoregressive OTU–Taxa pretraining

The model is pretrained autoregressively using a Transformer architecture [32]. The model receives a beginning-of-sample (BOS) token followed by the abundance-ranked OTU–taxonomy pairs and is trained to predict the next OTU and its taxonomy from the previous community context. This objective defines a generative model over the complete abundance-ranked microbiome profiles. The paired-token architecture and autoregressive pretraining setup are shown in Additional file 1: Supplementary Figure S1.

At each abundance-rank position, the input representation sums learned OTU, taxonomy, and rank embeddings. With embedding matrices *E*_otu_, *E*_tax_, and *E*_rank_, the representation at position *i* is

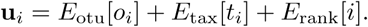

Raw counts and binned abundance values are not model inputs. A causally masked Transformer maps these representations to contextual states and uses only the preceding paired tokens to predict the next OTU and its assigned taxonomy.

For contextual state **h***_i_*, let **x***_<i_* denote the paired-token prefix strictly before position *i*, and let *θ* denote the model parameters. The two heads reuse their input embeddings as output projections, giving *P_θ_*(*o* | **x***_<i_*) = softmax(*E*_otu_**h***_i_*)*_o_* and *P_θ_*(*t* | **x***_<i_*) = softmax(*E*_tax_**h***_i_*)*_t_*, where *o* ∈ V_otu_ and *t* ∈ V_tax_ index candidate components of the corresponding softmax outputs. Taxonomy supervision follows the assigned path 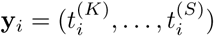 over ranks R = (*K, P, C, O, F, G, S*), denoting kingdom, phylum, class, order, family, genus, and species. Its hierarchy-consistent likelihood factorizes as

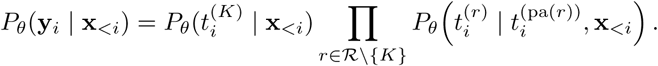

Here, pa(*r*) is the immediately preceding rank. The kingdom probability is normalized over kingdom nodes; each lower-rank probability is normalized over children of the assigned parent. When a rank is unresolved, its rank-specific unknown child is used and deeper ranks are excluded, preserving a truncated, valid path. The next-OTU loss *L_otu_* and taxonomy-path loss 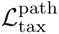 average their negative log-likelihoods over non-padding supervised positions. Complete notation and decoding are given in Additional file 1: Supplementary Methods M1C.

The joint pretraining objective is

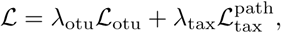

with *λ*_otu_ = *λ*_tax_ = 1 for the reported model. Padding and undefined taxonomy labels, including the taxonomy target at the terminal end-of-sample (EOS) position, are excluded.

The final model uses embedding dimension *d* = 256, six Transformer layers, four attention heads, feed-forward dimension 1024, *L* = 500, learned rank embeddings, and tied input/output projections. The MAP split contained 1,796,250 training, 20,000 validation, and 20,000 held-out samples; we selected the checkpoint with the lowest combined validation loss. Complete architecture and training settings are given in Additional file 1: Supplementary Methods M1B and M1D, the optimization history in Additional file 1: Supplementary Figure S2, and downstream settings in Additional file 1: Supplementary Methods M2B.

### 2.3 Sample classification

For the MAP-derived task panel, sample classification evaluates the mean-pooled pretrained representation for prediction of biological and environmental metadata, including body site, host phenotype, environment, disease status, and experimental condition. We compare this representation with supervised sample-by-OTU abundance classifiers.

We constructed 639 MAP-derived tasks, each confined to one study accession. After support and class-balance screening, two large language model (LLM)-assisted passes selected biologically or environmentally meaningful fields and removed constant, missing-dominated, protocol-related, or redundant ones using only field names and class-value counts. The frozen manifest and full curation protocol are described in Additional file 1: Supplementary Methods M2A. Of the 303,124 unique outer-test profile identifiers in the 639-task panel, 296,412 (97.79%) occurred in the self-supervised training partition. In the primary 250-task subset, the corresponding count was 104,947 of 107,327 (97.78%). All 404 study accessions in the full panel and all 137 in the primary subset were represented in that training partition; the remaining evaluation profiles belonged to the pretraining validation or test partitions. These benchmarks therefore assess in-domain prediction of withheld metadata labels. Task labels were reserved for downstream cross-validation [25, 26]. The panel contained 449 binary and 190 multiclass tasks, with a median of 593 samples per task (range 208–6,837) and 2–18 classes.

Disjoint partitions were feasible for 256 tasks. The primary benchmark comprises 250 of the 639 tasks with complete input coverage for all six methods and valid subjector experimental-unit-disjoint validation splits. Samples from the same subject or experimental unit are kept together in each split. The remaining 389 tasks use stratified sample-level folds and are reported separately in Additional file 1: Supplementary Table S5 and Figure S3. All fitting and model selection remained within each outer-training partition; Additional file 1: Supplementary Methods M2B gives the complete protocol.

Within each outer fold, we reload and fine-tune the pretrained encoder. For contextual states **h***_i_* and padding indicators *m_i_*, the sample embedding is ***Z****_x_* = (Σ*m_i_h_i_*)/(*Σ_i_m_i_*) A classification head is trained with class-weighted cross-entropy using only training-fold labels.

We compare OTU–Taxa with the MAP-retrained Pope-MAP, MGM, and Way-point models. Pope-MAP uses abundance-ranked OTUs, whereas MGM and Waypoint use abundance-ordered taxonomy profiles. All four pretrained models therefore use MAP-derived source profiles before downstream evaluation. Logistic Regression and class-balanced Random Forest use task-specific OTU-abundance features under the same outer cross-validation protocol. Adapter and training details are given in Additional file 1: Supplementary Methods M2B.

To assess the contribution of taxonomy inputs and supervision to classification, we additionally pretrained three matched conditions: paired OTU–taxonomy inputs with the joint objective, paired inputs with OTU loss only, and OTU-only inputs with OTU loss only. These conditions shared initialization, corpus partitions, backbone, and training budget, and were fine-tuned using identical classification folds and settings. The complete control protocol is given in Additional file 1: Supplementary Methods M1F.

For each task, we average balanced accuracy and the areas under the receiver operating characteristic (AUROC) and precision–recall (AUPR) curves across folds; multiclass AUROC and AUPR use macro one-vs-rest summaries. Metric-specific Fried-man tests precede Nemenyi comparisons at *α* = 0.05, with Holm adjustment across the three omnibus tests [33]. A critical-difference connection indicates that the comparison did not detect a rank difference at the selected threshold and is not an equivalence test. We also repeat the omnibus analysis after averaging task ranks within study accession. Additional file 1: Supplementary Methods M2C gives the ranking details.

### 2.4 Sample generation and disease-ecology readouts

The autoregressive objective makes OTU–Taxa a generative model of microbial communities: given the abundance-ordered prefix of a sample, the model assigns separate probability distributions to the next OTU and its taxonomy label. The analyses below use the OTU distribution; generated OTUs are paired with their fixed taxonomy annotations. We use this distribution for unconditional generation, conditional community completion, and exploratory disease-ecology readouts.

For unconditional generation, across ten fixed generation seeds, we sampled 3,000 complete communities per seed from the pretrained autoregressive model and compared them with 5,000 held-out test communities. In MAP abundance profiles, reads assigned to the same OTU are aggregated into a single count, so each sample contains one entry per distinct OTU. Generation preserves this structure through a no-repeat sampling constraint. Generation begins with the beginning-of-sample token. At step *i*, previously generated OTUs are paired with their fixed taxonomy annotations to form **x***_<i_*, and the model provides the next-OTU probability distribution *P_θ_*(*o* | **x***_<i_*). Probabilities assigned to previously emitted OTUs are set to zero, and the remaining distribution is renormalized before sampling. Generation stops at the end-of-sample token or maximum length. Empirical retrieval references sampled training profiles either globally or from the same study as the paired held-out target. We report OTU prevalence and top-200 OTU co-occurrence fidelity; full definitions and uncertainty estimates are provided in Additional file 1: Supplementary Methods M3A and Additional file 1: Supplementary Table S7.

Community completion asks whether the model can recover withheld lower-abundance members after observing the most-abundant half of the same sample. For each held-out sample with *L_x_* ≥ 8 OTUs, let *h* = ⌊*L_x_/*2⌋ be the number of observed prefix OTUs. Each candidate *o* receives the autoregressive (AR) score *s*_AR_(*o*) = *P_θ_*(*o* | BOS*, o*_1_*, . . . , o_h_*), with observed OTUs removed before ranking. The final hierarchy-aware checkpoint was evaluated on 2,000 held-out test samples. Base-lines were global popularity, word2vec [34], learned-embedding co-occurrence, and MAP-training-set nearest-sample retrieval (Additional file 1: Supplementary Methods M3A and Additional file 1: Supplementary Table S7). This is a one-step conditional ranking: all withheld OTUs are evaluated against the same fixed observed prefix, which is not extended by model predictions. We report Recall@*k* (recall among the top *k* candidates), NDCG@*k* (normalized discounted cumulative gain), and AUPRC (area under the precision–recall curve) for *k* ∈ {10, 50, 100}.

The main completion results use fixed-prefix ranking; alternative prediction procedures are compared in Additional file 1: Supplementary Methods M3A and Supplementary Results R4.

We next explored whether the model’s generative probabilities reflect population-associated genus abundance shifts across disease and ecological contrasts. This analysis assesses whether genera with higher observed abundance in one population also receive earlier model-expected abundance ranks in that population. OTU-level probabilities are collapsed to genus *g* by summing over assigned OTUs, 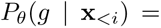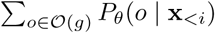, where O(*g*) is the set of OTU tokens assigned to genus *g*.

For IBD analyses, model-derived genus scores are compared with the signed pooled direction of curated BugSigDB signatures [35]; positive values indicate more reports of increased than decreased abundance. The snapshot, filtering, genus collapse, and signature weighting are detailed in Additional file 1: Supplementary Methods M3B.

For the within-cohort analysis, we used 100 prespecified binary study–metadata-field tasks. In each task, the two selected values of one metadata field divided samples from the same study into groups denoted neg and pos to orient the comparison. Because the autoregressive sequence is abundance ordered, we summarize where the model assigns probability to genus *g* separately for each group *c* ∈ {neg, pos} using the probability-weighted position

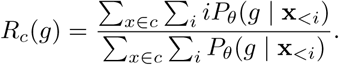

The heuristic model abundance-rank score is *R*_neg_(*g*)−*R*_pos_(*g*); positive values indicate that the model assigns relatively more probability to genus *g* at earlier positions in the pos group. The score is measured in abundance-rank positions; it is not a predicted relative abundance or a calibrated fold change.

For within-cohort comparisons, OTU relative abundances were summed by genus and used to calculate the observed genus-level log fold change (logFC) between the two groups. Within each task, labels were permuted 10,000 times while respecting validated subject or repeated-unit identifiers where available; sample-level permutations preserving group sizes were used when no validated identifier was available. Within each task, Benjamini–Hochberg (BH) adjustment controlled the false-discovery rate (FDR) across genus-level permutation *p*-values during genus selection. Of the 100 tasks, 78 had at least 10 genera with significant abundance differences (*q* ≤ 0.05, where *q* is the BH-adjusted permutation *p*-value) and included genera enriched in both groups, allowing AUROC to be calculated. Within each retained task, AUROC measured whether the model abundance-rank score distinguished genera enriched in the pos group from those enriched in the neg group, while Spearman correlation compared the model score with the signed logFC. This BH correction is distinct from Holm adjustment across the three classification omnibus tests. For computational tractability, model scores used at most 400 profiles per class and teacher-forced prefixes through position 300, whereas the count analysis used every eligible profile. Full construction and subset definitions are provided in Additional file 1: Supplementary Methods M3C.

### 2.5 Context-based taxonomy-assignment recovery

The context-based taxonomy-assignment recovery task asks whether the surrounding microbial community contains useful information for recovering SINTAX/QIIME2 assignments of OTUs represented in the pretrained vocabulary as target-sequence evidence is reduced. Sequence-based classifiers such as SINTAX [30], Quantitative Insights Into Microbial Ecology 2 (QIIME2) [14, 36], and RAxTax [37] assign taxonomy from the representative OTU sequence. In contrast, OTU–Taxa predicts taxonomy from the sample prefix before the target OTU, and the taxonomy head is decoded as a hierarchy-consistent path as described in Additional file 1: Supplementary Methods M1C. Prior work shows that environment-specific abundance priors can improve sequence-taxonomy accuracy, whereas algorithmic changes alone offer limited gains when short sequences remain ambiguous [38, 39].

For the main comparison, we use SINTAX-derived labels, matching the label source used during OTU–Taxa pretraining. The paired SINTAX-aligned comparison includes 5,666 OTUs with eligible full-sequence labels and held-out MAP samples containing them for context-only evaluation. Each representative 16S sequence is degraded by random substitutions, random deletions, or centered truncation, then reclassified and scored against its full-sequence assignment. These controlled stress tests create a sequence-evidence gradient inspired by base-calling errors, local sequence disruption, and marker-region or length effects [12, 36, 40]; the strongest levels are extreme cases rather than typical clean-read error rates. QIIME2 provides a second taxonomy-label source for a matched OTU–Taxa experiment. RAxTax is included as an additional sequence-classifier robustness reference and is evaluated on the same OTUs and SIN-TAX reference assignments as the SINTAX-aligned comparison. Exact degradation grids and scoring details are provided in Additional file 1: Supplementary Methods M4A. In the QIIME2 sequence-truncation analysis, a fixed naive Bayes classifier classified progressively shortened queries; the resulting curve shows how its predictions change as query length decreases.

For OTU–Taxa, each selected OTU is evaluated in held-out samples that contain that OTU. At each occurrence of that OTU in a held-out sample, at position *j*, the model receives only the observed prefix before *j*; the target OTU identity and taxonomy are withheld. Because sequence-based classifiers produce one assignment per representative sequence, we choose, for each OTU and rank, the taxonomy label predicted most often across its evaluated held-out sample occurrences.

Taxonomy labels form paths through a hierarchy, so exact-match accuracy alone is too coarse: a species-level mistake with the correct genus is less severe than an error beginning at phylum. We therefore report per-rank agreement with classifier assignments and hierarchical rank accuracy (rankACC). For each OTU, rankACC is the fraction of available reference ranks recovered consecutively in taxonomic order before the first disagreement. Ranks without a reference annotation are excluded from scoring. It ranges from 0 to 1 and gives partial credit when an error occurs only at a lower rank. Full mathematical definition, per-rank agreement, and complete-path exact-match rates are provided in Additional file 1: Supplementary Methods M4A and Additional file 1: Supplementary Table S9. Here, “reference” denotes the classifier assignment obtained from the intact representative sequence. The benchmark therefore measures recovery of, or agreement with, that label path rather than accuracy against independently curated taxonomic truth.

The final comparison reports sequence-degradation curves together with the corresponding context-only OTU–Taxa score. SINTAX and RAxTax use the same SINTAX reference assignments and SINTAX-trained context-only score, whereas QIIME2 uses its matched QIIME2 labels and context-only model. The sequence and context scores are evaluated separately; this experiment does not train or test a hybrid classifier that combines both evidence sources.

## 3 Results

### 3.1 Sample classification across MAP-derived tasks

We first evaluated two recognizable MAP metadata tasks, American Gut Project (AGP) IBD status [41] and Human Microbiome Project (HMP) body site [42], as compact references before aggregating across the broader panel. Under subject-disjoint validation, MGM had the highest mean values for all three metrics on AGP IBD, although its AUPR varied substantially across folds [mean (standard deviation, SD), 0.181 (0.193)]; on HMP body site, OTU–Taxa had the highest balanced accuracy and AUPR, while Pope-MAP had the highest mean AUROC (Additional file 1: Supplementary Table S6).

Method ranks differed across the 250 disjoint-fold tasks for all three metrics (Holm-adjusted Friedman *p <* 2 × 10*^−^*^65^; Figure 1). The omnibus conclusions remained significant after ranks were first averaged within the 137 study accessions (Holm-adjusted *p <* 2 × 10*^−^*^47^), reducing the influence of accessions that contributed multiple tasks. Across-task mean AUROC (SD) was 0.749 (0.191), 0.744 (0.189), 0.730 (0.192), 0.725 (0.191), 0.707 (0.179), and 0.660 (0.172) for Random Forest, OTU–Taxa, MGM, Waypoint, Logistic Regression, and Pope-MAP, respectively. Random Forest and OTU–Taxa tied in balanced accuracy [0.674 (0.188) and 0.674 (0.190)], and Random Forest led AUPR [0.705 (0.231) versus 0.695 (0.229)] (Additional file 1: Supplementary Table S4A). Giving each study accession equal weight generally yielded slightly higher mean AUROC and AUPR, while preserving the ordering of methods for these metrics. The aggregate model comparison was therefore little affected by unequal numbers of tasks per accession (Additional file 1: Figure S4). For every metric, the Nemenyi comparisons detected neither a rank difference between MGM and Waypoint nor one between OTU–Taxa and Random Forest (Figure 1). OTU–Taxa was therefore the highest-ranked pretrained model. On the 389 stratified sample-level tasks, OTU– Taxa again led the pretrained models, but Random Forest led all three metrics by a wider margin (mean AUROC (SD), 0.886 (0.145) versus 0.871 (0.154); Additional file 1: Supplementary Table S5 and Figure S3). The pretrained model provides a reusable sample-level signal, while competitive OTU-abundance classifiers remain strong. In the matched-pretraining comparison across the same 250 tasks, the three conditions showed no statistically significant classification differences after Holm adjustment (Additional file 1: Supplementary Results R3, Table S4B–C and Figure S5).

**Fig. 1.**
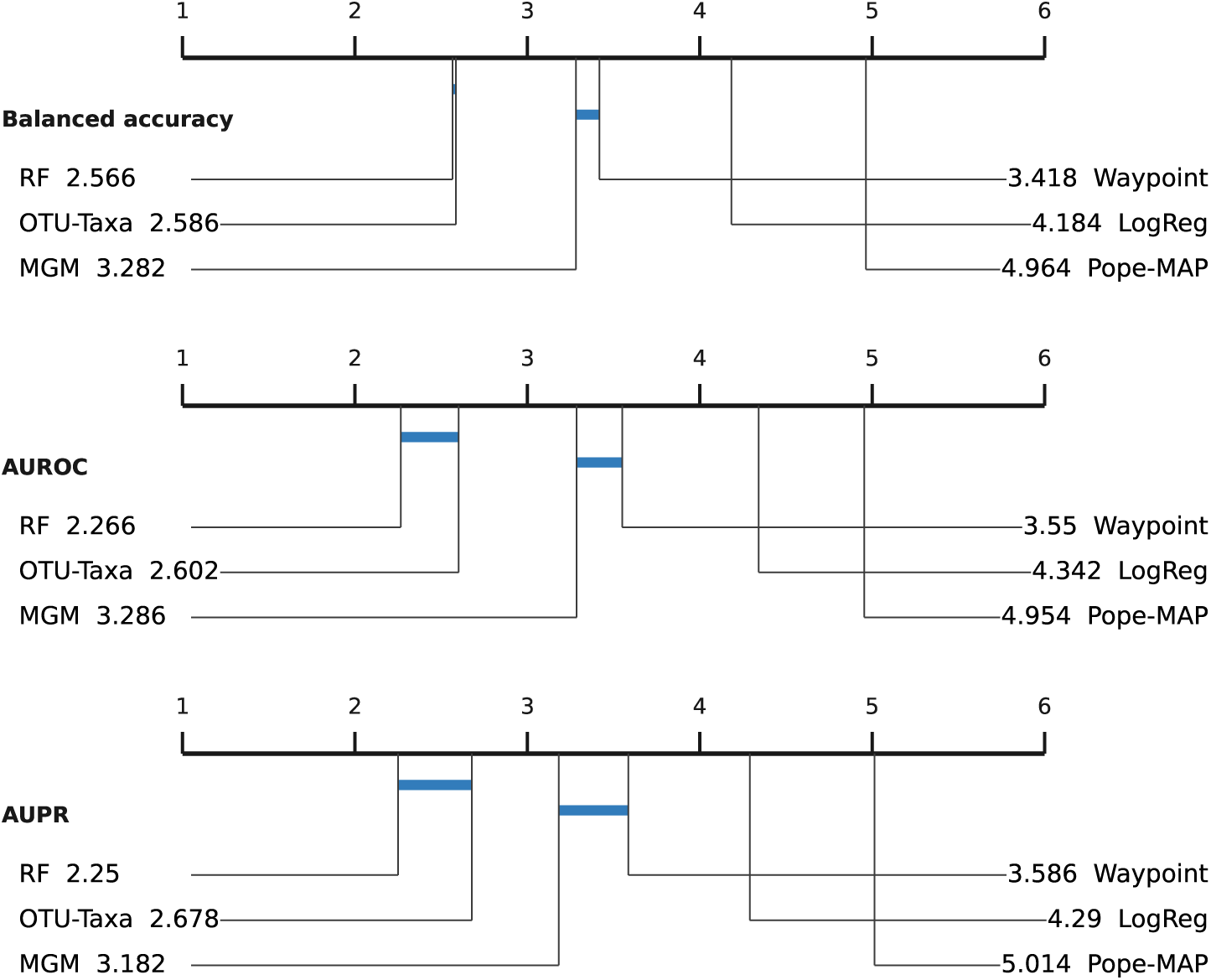
Critical-difference summaries for 250 classification tasks evaluated with disjoint folds. Lower average rank is better; the Nemenyi critical difference is 0.48. Blue segments connect methods for which the Nemenyi analysis did not detect a rank difference of at least this value; a connection does not establish equivalent performance.

### 3.2 Sample generation and disease-ecology readouts

To test whether the autoregressive model produces community-level structure, we compared generated samples with held-out MAP communities and empirical training profiles. Across ten generation seeds, mean (SD) top-200 OTU co-occurrence Pearson correlation was 0.9771 (0.0010) for OTU–Taxa, 0.9854 (0.0009) for global training-profile retrieval, and 0.9912 (0.0005) for same-study retrieval. Same-study retrieval was available for 2,904 of the 3,000 targets per replicate. Generated output was not dominated by exact copies: only 19 of 30,000 profiles exactly matched a training sample. Nearest-training OTU-set Jaccard similarity had a median of 0.249 and a 99th percentile of 0.625 (Additional file 1: Supplementary Results R4).

Conditional completion gave a second view of the same behavior. Under the canonical MAP tie order, sample retrieval gave stronger early ranking (NDCG@10, 0.482 versus 0.389; paired OTU–Taxa-minus-retrieval difference, −0.093 [95% profilebootstrap confidence interval (CI), −0.107, −0.079]) and higher AUPRC (0.218 versus 0.207), whereas OTU–Taxa recovered a broader fraction of withheld members at larger cutoffs (recall@100, 0.393 versus 0.322; paired difference, 0.071 [0.058, 0.083]). In an audit of 1,000 held-out profiles, 97.1% contained at least one exact-count tie. Tied blocks contained 75.3% of model-visible OTUs, increasing to 97.5% at ranks 101– 499, where within-block positions followed MAP record order. Permuting equal-count OTUs changed the next-token distribution (mean Jensen–Shannon divergence 0.286; 95% CI, 0.272–0.301). In a separate evaluation of 2,000 completion queries, averaging over five alternative within-prefix tie orderings, all seven mean completion metrics increased. NDCG@100 increased from 0.371 to 0.408 (change, 0.036 [95% CI, 0.034, 0.039]; retrieval, 0.390), and AUPRC increased from 0.207 to 0.239 (change, 0.032 [0.030, 0.034]; retrieval, 0.218). The broader pattern of stronger retrieval at early ranks and broader OTU–Taxa recovery was preserved. Full uncertainty estimates and the cooccurrence and popularity baselines are provided in Additional file 1: Supplementary Tables S7 and S3B.

We explored model–count agreement among genera with significant abundance differences (*q* ≤ 0.05), retaining contrasts with at least 10 selected genera as defined in Section 2.4. Within each contrast, the count data determined both genus selection and the reference enrichment/depletion directions. The reported AUROC therefore quantifies genus-level direction agreement conditional on this selected set. In a plant drought experiment, the model abundance-rank contrast agreed strongly with the count-logFC contrast across 170 significant genera (AUROC 0.828; Spearman 0.495; Additional file 1: Supplementary Figure S7). In contrast, a lake-water versus stream-water comparison with 28 significant genera was systematically inverted rather than merely uninformative (AUROC 0.115; Spearman −0.294). These examples illustrate that the readout is a task-dependent abundance-rank signal: the same pretrained model can align strongly with a population contrast in one setting and invert it in another, so single-task readouts need the count-based comparison alongside them.

Across these 78 contrasts, agreement increased when the analysis was restricted to more abundant genera (Figure 2). The conditional mean AUROC across all selected genera was 0.588 (95% study-cluster bootstrap interval 0.536–0.642), exceeding the within-contrast permutation null (null mean 0.4998; one-sided *p <* 0.001). Mean AUROC increased from 0.588 across all FDR-selected genera to 0.673 and 0.718 when retaining selected genera at or above the 50th and 75th abundance percentiles, respectively. Percentiles were calculated among all genera shared by the count and model tables within each contrast, using pooled mean relative abundance. Within each contrast, the corresponding increases relative to the AUROC calculated using all selected genera were 0.084 (95% study-accession bootstrap CI, 0.068–0.101) and 0.129 (0.106– 0.154), respectively. Agreement was therefore concentrated among more abundant genera, consistent with more stable rank information in the abundance-ordered head of the profile and sparser observations in the low-abundance tail (Additional file 1: Supplementary Figure S8).

**Fig. 2.**
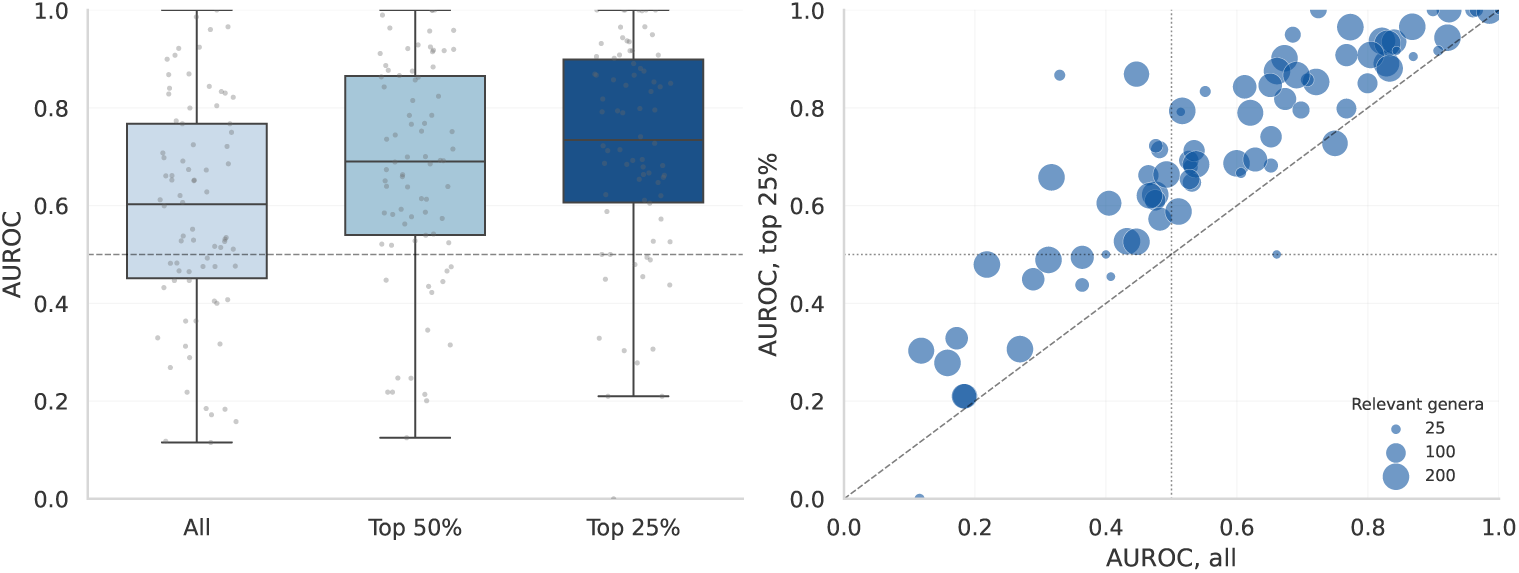
Agreement between model abundance-rank and observed count-logFC directions. The left panel shows the conditional AUROC distribution across contrasts for all task-specific FDR-selected genera and for subsets retaining selected genera at or above the 50th and 75th abundance percentiles among all shared genera. Abundance is measured by pooled mean relative abundance within each contrast; grey points are individual contrasts and the dashed line marks chance (0.5). The right panel compares each contrast’s top-quarter AUROC with its all-selected-genera AUROC. Marker area represents the number of selected genera, the diagonal is equality, and dotted lines mark AUROC 0.5.

For IBD, we compared the abundance-rank readout with pooled BugSigDB directions as a descriptive external reference. Concordance AUROCs for the model-derived score and observed count-logFC were 0.674 and 0.406, respectively, across 122 genera in AGP [41], and 0.770 and 0.808 across 72 genera in the Halfvarson cohort [7]. The cohorts behave differently: in Halfvarson the model tracks the literature directions that the counts themselves recover, whereas in AGP the model agrees with BugSigDB more closely than the AGP counts do, which are anti-concordant. We there-fore read the AGP value as consistent with broad cross-study gut associations carried through pretraining rather than as an AGP-specific disease contrast (Additional file 1: Supplementary Table S8).

### 3.3 Context-based taxonomy-assignment recovery

To test whether sample context supports taxonomy-assignment recovery, we first evaluated context-only predictions for agreement with assignments in the SINTAX label space. Mean hierarchical rank accuracy relative to the SINTAX assignments reached 0.456 (95% CI, 0.448–0.464) across 5,666 OTUs (Additional file 1: Supplementary Figure S9). Agreement was concentrated at higher ranks: per-rank agreement was 0.991 at kingdom, 0.602 at phylum, 0.158 at genus, and 0.009 at species, and 0.101 of complete paths matched (Additional file 1: Supplementary Table S9). Intact sequences remained the strongest evidence, but agreement with the corresponding full-sequence assignment declined under controlled degradation while the context-only score provided a target-sequence-independent reference from the surrounding community (Figure 3). The crossing curves identify degradation levels at which the separate context score exceeds classifier self-agreement; they do not demonstrate that combining sequence and context improves an assignment.

**Fig. 3.**
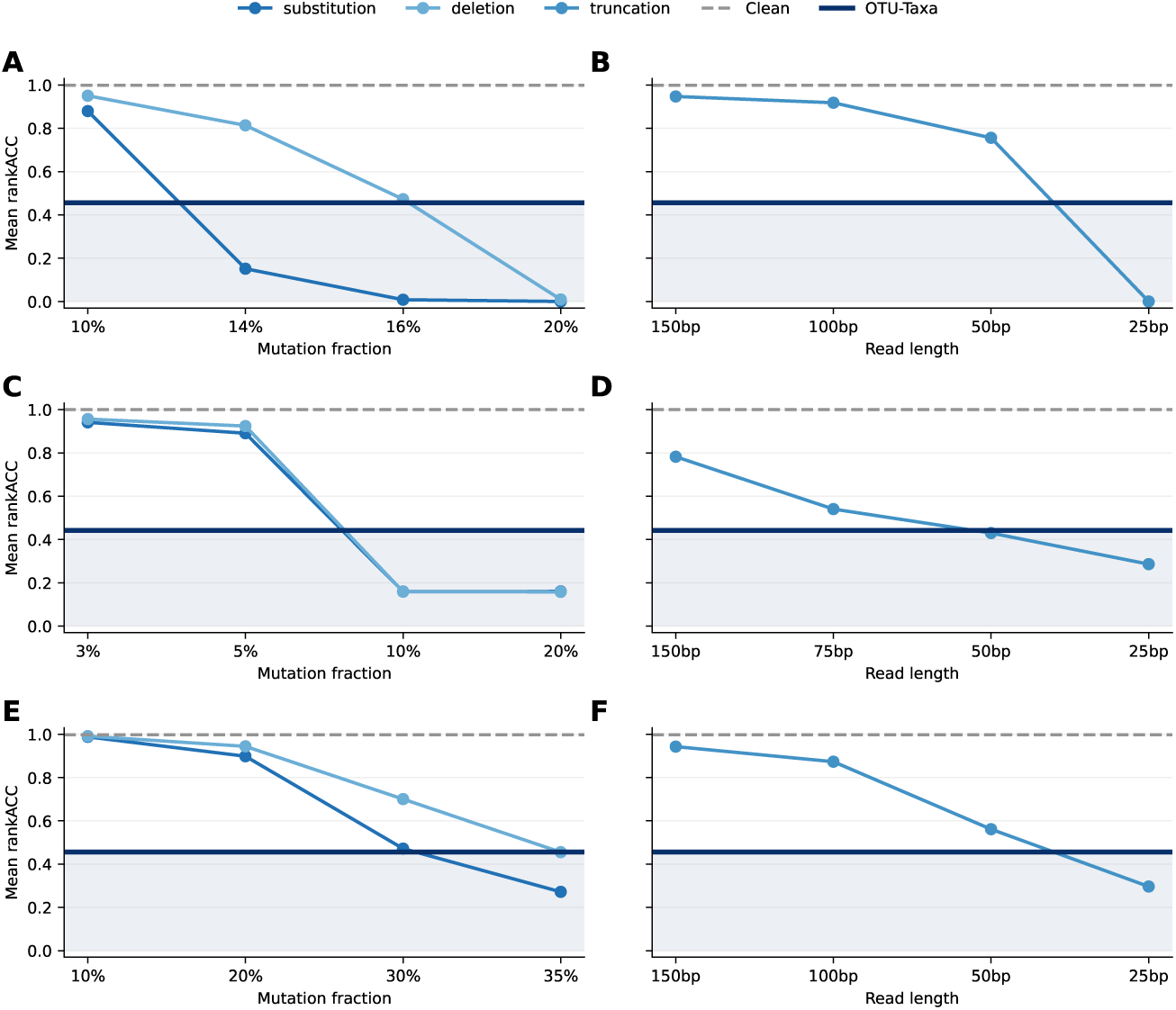
Sequence-degradation curves and context-based taxonomy-assignment recovery for SINTAX (A,B), QIIME2 (C,D), and RAxTax (E,F). Panels A, C, and E show random substitutions and deletions; panels B, D, and F show centered truncation. SINTAX and RAxTax predictions are scored against intact-sequence SINTAX assignments, whereas QIIME2 predictions are scored against intact-sequence QIIME2 assignments. Grey dashed lines show agreement of intact-sequence predictions with these references. Dark horizontal lines show the SINTAX-trained context-only OTU–Taxa score in panels A, B, E, and F, and the QIIME2-trained score in panels C and D.

The sample-context ablations in Additional file 1: Supplementary Figure S10 separate the sample-specific signal from simpler global-frequency explanations. The full OTU–taxonomy context gave the highest SINTAX-aligned rankACC among the ablations (0.456), compared with 0.310 for a global-prior predictor and 0.260 for shuffled sample context; OTU-only and taxonomy-only contexts retained intermediate signal (0.433 and 0.423). A visible-context fraction analysis further shows how the available prefix context affects the readout (Additional file 1: Supplementary Figure S11). The matched QIIME2 model showed a similar rank-wise error pattern (Additional file 1: Supplementary Figure S12). For this secondary label source, the context-only OTU– Taxa evaluation reached mean rank accuracy relative to QIIME2 assignments of 0.442 (95% CI, 0.433–0.451) across 4,168 OTUs.

This controlled benchmark shows that the model can recover part of an intact-sequence classifier’s label path from local community context when the target identity is withheld. The scores quantify classifier-assignment recovery for known-vocabulary OTUs with observed community context; performance when sequence evidence is degraded across an entire experiment remains untested.

## 4 Discussion

The paired OTU–Taxa model learned a reusable representation of abundance-ranked 16S community profiles that supported sample classification, community generation and completion, disease/ecology readouts, and recovery of taxonomic labels from sample context. It was the highest-ranked pretrained method in the primary classification benchmark, while Random Forest had slightly higher mean AUROC and AUPR. The two methods belonged to the same Nemenyi-connected group, meaning that the rank comparison did not resolve a difference at the chosen threshold; this result is not evidence of equivalence. The omnibus result remained stable after ranks were averaged within study accession, although the panel combines heterogeneous biological, environmental, and technical labels. The different leaders on AGP IBD and HMP body site further show that aggregate ranks should inform, rather than replace, task-specific method choice. The pretrained comparators also retain model-specific vocabularies, capacities, and MAP adaptations, so their ranks compare complete systems rather than isolate an architecture or representation effect.

The architecture couples fine-grained OTU identity, a hierarchy-structured taxonomy annotation, and abundance rank in one pretrained representation. The experiments therefore evaluate this bundle. The OTU-only and taxonomy-only context ablations remove one input channel from the already trained paired model and show that both channels retain information, but they do not estimate the causal contribution of taxonomy to classification or generation. To examine its contribution to classification, we additionally compared matched pretraining conditions with and without taxonomy inputs and supervision, finding no significant differences in fine-tuned classification performance (Additional file 1: Supplementary Results R3, Table S4B–C and Figure S5). One possible explanation is that OTU identities already provide much of the information needed for these classification tasks, making the associated taxonomy annotations partly redundant. This finding applies to the classification tasks evaluated here; the aim of the joint representation is to support both OTU-level and hierarchy-structured taxonomic representations within one model. The present evidence supports taxonomy awareness and use of both channels, while the benefit of adding taxonomy inputs and supervision for generation and context-based taxonomy-assignment recovery remains to be tested.

Generation and retrieval expose a useful distinction between learned and empirical community models. Retrieval better preserved held-out richness, prevalence, co-occurrence, and early completion ranks, while OTU–Taxa produced few exact training-set copies and recovered more withheld members at larger cutoffs. The 19 exact matches rule out pervasive exact duplication but do not exclude more diffuse forms of memorization. Completion also ranks every withheld member from one next-token distribution rather than rolling the model forward, so it tests one-step community association, not reconstruction of the entire missing suffix. Finally, abundance determines the ordering between different count levels, while positions within equalcount blocks follow a secondary ordering convention. Exact ties contained 75.3% of visible OTUs, and alternative within-tie orders improved all seven completion met-rics. The broad early-ranking versus wider-recall pattern was stable. Fine-grained autoregressive scores therefore depend partly on that convention, particularly in the low-abundance tail. Randomizing ties during training and reporting performance averaged over valid orders, or using set-aware architectures, would make this uncertainty part of the model rather than a post hoc sensitivity analysis.

The disease-ecology analysis connected teacher-forced next-OTU probabilities with observed genus-level contrasts and found the strongest agreement among more abundant selected genera. The resulting AUROC is conditional on genera first selected by the count analysis, and the score is expressed in abundance-rank positions rather than relative abundance or fold change. Stronger agreement among abundant genera may reflect more stable biological signal, but it may also follow from more stable counts and from training an abundance-ordered next-token objective. The pooled BugSigDB comparison provides a complementary literature-facing readout, although it combines heterogeneous disease-versus-control and within-IBD signatures and weights signatures rather than studies. These exploratory, hypothesis-generating analyses assess agreement between the model readout and observed genus abundance shifts. They do not establish validated biomarkers, disease-prediction performance in new patients, or causal microbial responses.

The taxonomy-assignment recovery experiment quantified how much of the SIN-TAX or QIIME2 label path could be recovered from community context after the target OTU identity was withheld. Agreement was strongest at high taxonomic ranks, with SINTAX-aligned per-rank agreement declining from 0.602 at phylum to 0.158 at genus and 0.009 at species. The global-prior and shuffled-context controls show that the full score contains sample-specific information, while the OTU-only and taxonomyonly controls show that neither channel is solely responsible. These values measure agreement with intact-sequence classifier assignments rather than taxonomic truth. Intact sequences remained the strongest evidence, and the current study compares sequence and context rather than combining them. A direct next step is a hybrid classifier in which sequence likelihood and community context are integrated and calibrated, followed by a direct comparison with its sequence-only counterpart under the same label definition. The target OTUs already belong to the pretrained vocabulary and are evaluated in held-out sample contexts. In addition, each random corruption condition uses one deterministic realization; the curves do not quantify perturbation-seed variability. The strongest degradation levels should remain stress tests, not proxies for typical clean-read error rates.

The released checkpoint provides a reusable representation within the MAP 97% closed-reference 16S vocabulary. New profiles must first be mapped to these identifiers because the model has no learned representation for an unmatched ASV or OTU. Classification uses fold-local labels and held-out outer folds. Most evaluated profiles occurred in the self-supervised training partition, so the MAP task panel tests indomain prediction of withheld metadata labels.

## 5 Conclusions

Within the MAP 97% closed-reference vocabulary, the pretrained representation supports in-domain prediction of metadata labels alongside generative and taxonomic readouts. The results also identify a practical division of labour: supervised abundance models, empirical retrieval, and intact-sequence classifiers remain stronger choices for several corresponding endpoints, while OTU–Taxa provides one shared contextual representation across all four analyses.

## Supporting information

aditional file

## Declarations

### Ethics approval and consent to participate

Not applicable. This study involved the secondary analysis of publicly available, de-identified microbiome profiles and associated non-identifying metadata. No participants were recruited, no new biological samples were collected, and no identifiable personal data were accessed.

### Consent for publication

Not applicable.

### Availability of data and materials

The OTU–Taxa source code is publicly available in the github.com/ bio-ontology-research-group/otu-taxa-autoregressive under the MIT License [43]. The released workflows use Python 3.9 or later and PyTorch, were tested on Linux, and require a CUDA-compatible PyTorch installation for GPU execution. The MAP-retrained MGM, Waypoint, and PopeMAP workflows, configurations, and evaluation code are available in the github.com/bio-ontology-research-group/microbiome foundation model benchmarks [44]. The final OTU–Taxa checkpoint, vocabularies, taxonomy hierarchy, and inference example are available from the huggingface.co/bio-ontology-research-group/otu-taxa [45]. Processed inputs, the frozen 639-task benchmark and fold assignments, model predictions, result tables, and figure-source data are available from the huggingface.co/datasets/bio-ontology-research-group/otu-taxa-paper-artifacts [46]. Raw Microbe Atlas Project source files are not redistributed; their use remains subject to the conditions of the original source. The released resources are publicly accessible. The OTU–Taxa repository contains code for pretraining and each analysis, together with results notebooks organized by the paper’s classification, generation/completion and disease-ecology, and context-based taxonomy-assignment recovery sections. Other notebooks cover pretraining diagnostics and abundance-order sensitivity. A section-to-notebook guide identifies the code and released inputs used for each analysis (https://github.com/bio-ontology-research-group/otu-taxa-autoregressive/tree/ main/results notebooks).

### Authors’ contributions

Conceptualization was carried out by R.He. and R.Ho. Methodology was developed by R.He., R.Ho., and H.P. H.P. implemented the models and performed data curation, with additional contributions from R.He. Formal analysis was conducted by H.P., R.He., and R.Ho. H.P. wrote the original manuscript draft, and all authors contributed to writing, reviewing, and editing the final manuscript. Supervision was provided by R.He. and R.Ho. All authors read and approved the final manuscript.

## Acknowledgements

Not applicable.

## Additional files

**Additional file 1.** Filename: Additional file 1.pdf. Format: PDF. Title: Supplementary methods and results. Description: extended pretraining and representation methods, classification protocols, generation and disease-ecology analyses, context-based taxonomy-assignment recovery methods, and the corresponding supplementary tables and figures.

