## Supplementary material for "OTU–Taxa: a taxonomy-aware pretrained model for 16S microbiome profiles": aditional file

### Additional file 1

#### Supplementary Methods

In prose, dispersion summaries are reported as mean (standard deviation, SD); tables may use the equivalent mean ± SD notation. SD denotes the sample standard deviation across the stated replication unit: outer folds for the two reference classification tasks, task-level cross-validation means for the 250- and 389-task summaries, and stochastic replicates for unconditional generation. Confidence intervals (CIs) are used for aggregate estimates computed from many evaluation units: completion metrics use profile-level bootstrap intervals, disease-ecology summaries use study-accession bootstrap intervals, mean hierarchical rank accuracy (rankACC) uses a normal-approximation interval across evaluable OTUs, and binary per-rank and complete-path agreement proportions for taxonomy assignments use Wilson intervals. Thus, SD describes observed dispersion among repeated measurements, whereas a confidence interval describes uncertainty in the reported aggregate; the corresponding resampling procedures are specified with each analysis below.

##### Supplementary Methods M1. Pretraining and Sample Representation

###### M1A. Sample Filtering and Taxonomy Vocabularies

The analyzed profiles derive from 16S ribosomal RNA (rRNA) gene amplicon sequencing. The main Microbe Atlas Project (MAP) representation retains operational taxonomic units (OTUs) represented by at least four reads within each sample before abundance ranking. This read-count filter is applied independently per sample and does not change the global OTU vocabulary. Filtering very low-count amplicon features is a standard quality-control step for reducing likely sequencing or amplification noise, although exact thresholds vary across workflows [1, 2]. Observed OTUs are ordered by descending count using a stable sort. Equal-count OTUs therefore retain their order in the corresponding MAP sample record; this secondary order is inherited from the source profile rather than constructed from the OTU vocabulary index or randomized during tokenization. Taxonomy labels are assigned across the seven ranks $K, P, C, O, F, G, S$, corresponding to kingdom through species. A rank is accepted only when classifier confidence exceeds 80%; at the first rank that does not, that rank and all descendants are represented by rank-specific unknown nodes. Rare deeper assignments below that point are truncated, preserving a valid hierarchical path.

The tokenizer is indexed by the 62,200 identifiers in MAP's 97% closed-reference OTU system. This preserves the native feature system of the pretraining corpus rather than reconstructing an independent amplicon sequence variant (ASV) vocabulary. The released checkpoint has no generic unseen-OTU token. Applying it to another feature table therefore requires closed-reference mapping to these MAP identifiers; unmatched ASVs or OTUs are not represented directly.

Within MAP, each taxonomy token annotates its paired OTU with the deepest accepted node in the reference hierarchy. The paired representation therefore combines a fine-grained sequence-derived feature with its position in a shared biological hierarchy. Microbiome datasets can be distributed at different representation levels: reference-based feature tables preserve OTU or ASV identities, whereas other processed resources retain taxonomic profiles without the underlying feature assignments. OTU–Taxa retains aligned OTU and taxonomy channels so that community context can be expressed through either level within the learned vocabulary. The full-, OTU-only-, and taxonomy-only-context analyses characterize these channels within the MAP-derived representation. Because these are inference-time ablations of one paired checkpoint, they do not replace parameter-matched OTU-only or taxonomy-only pretraining controls.

##### M1B. Paired-Token Transformer Architecture

Let BOS and EOS denote the beginning- and end-of-sample tokens, respectively. For a sample $\mathbf{x}$, the OTU input sequence is $\mathbf{o}^{\text{in}} = (\text{BOS}, o_1, o_2, \ldots, o_{L_x})$, and the OTU target sequence is $\mathbf{o}^{\text{lab}} = (o_1, o_2, \ldots, o_{L_x}, \text{EOS})$. Analogously, the taxonomy input sequence is $\mathbf{t}^{\text{in}} = (\text{BOS}, t_1, t_2, \ldots, t_{L_x})$; taxonomy labels are assigned at positions whose targets are observed OTUs. At position $i$, the model predicts the next OTU and the taxonomy assigned to that OTU.

The model uses separate OTU and taxonomy embedding matrices, $E_{\text{otu}} \in \mathbb{R}^{|\mathcal{V}_{\text{otu}}| \times d}$ and $E_{\text{tax}} \in \mathbb{R}^{|\mathcal{V}_{\text{tax}}| \times d}$, where $d$ is the embedding dimension, and a learned abundance-rank embedding $E_{\text{rank}} \in \mathbb{R}^{L \times d}$. The beginning-of-sample token occupies position zero, and subsequent positions encode abundance rank. Raw counts and binned abundance values are not provided. The summed embeddings are layer-normalized, passed through embedding dropout, and truncated or padded to $L = 500$.

A Transformer encoder with a causal mask maps the input-embedding matrix $U = [\mathbf{u}_1; \ldots; \mathbf{u}_L]$ to contextual states $H = f_\theta(U) = [\mathbf{h}_1; \ldots; \mathbf{h}_L]$, where $\theta$ denotes the model parameters, so each prediction uses only the preceding abundance-ordered prefix. The OTU and taxonomy heads reuse their respective input embeddings as output projections, producing logits $\mathbf{z}_i^{\text{otu}} = E_{\text{otu}}\mathbf{h}_i$ and $\mathbf{z}_i^{\text{tax}} = E_{\text{tax}}\mathbf{h}_i$. Their softmax distributions define $P_\theta(o \mid \mathbf{x}_{<i})$ over OTU tokens and $P_\theta(t \mid \mathbf{x}_{<i})$ over taxonomy tokens, where $\mathbf{x}_{<i}$ is the paired-token prefix strictly before position $i$. Supplementary Methods M1C defines how the taxonomy distribution is trained and decoded as a valid hierarchical path.

##### M1C. Hierarchy-Aware Taxonomy Objective and Decoding

Let $\theta$ denote the model parameters and let $\mathcal{T} = (\mathcal{V}_{\text{tax}}, \mathcal{E})$ denote the directed, rank-ordered taxonomy hierarchy, where $\mathcal{V}_{\text{tax}}$ is the taxonomy-token vocabulary and $\mathcal{E}$ contains the permitted parent–child relations. The ordered rank set is $\mathcal{R} = (K, P, C, O, F, G, S)$, denoting kingdom, phylum, class, order, family, genus, and species. We use $\mathcal{V}_r \subset \mathcal{V}_{\text{tax}}$ for the tokens at rank $r$, including its unknown token, and $\mathcal{V}_{\text{otu}}$ for the OTU-token vocabulary. We use $\text{pa}(r)$ for the immediately preceding rank; for example, $\text{pa}(G) = F$.

For a sample $x$, $\mathbf{x}_{<i}$ is the sequence of paired OTU and taxonomy inputs strictly before position $i$. Let $\mathcal{I}_x^{\text{otu}}$ be the non-padding positions with a supervised OTU target, including the terminal EOS target, and let $\mathcal{I}_x^{\text{tax}}$ be the positions whose next observed OTU has a taxonomy target; padding and the terminal EOS position are excluded from the second set. At each $i \in \mathcal{I}_x^{\text{otu}}$, $o_i^{\text{lab}} \in \mathcal{V}_{\text{otu}}$ is the next-OTU target, with EOS used at the terminal position. For $i \in \mathcal{I}_x^{\text{tax}}$, the taxonomy assigned to that OTU defines the rank-wise target path

$$\mathbf{y}_i = (t_i^{(K)}, t_i^{(P)}, t_i^{(C)}, t_i^{(O)}, t_i^{(F)}, t_i^{(G)}, t_i^{(S)}),$$

where $t_i^{(r)} \in \mathcal{V}_r$ is the target taxonomy token at rank $r$.

The taxonomy head produces a logit $z_{i,v}^{\text{tax}}$ for every candidate $v \in \mathcal{V}_{\text{tax}}$. At the kingdom rank, the softmax is normalized over $\mathcal{V}_K$. At each lower rank $r$, it is normalized only over

$$\text{Child}_r(u) = \{v \in \mathcal{V}_r : (u \to v) \in \mathcal{E}\},$$

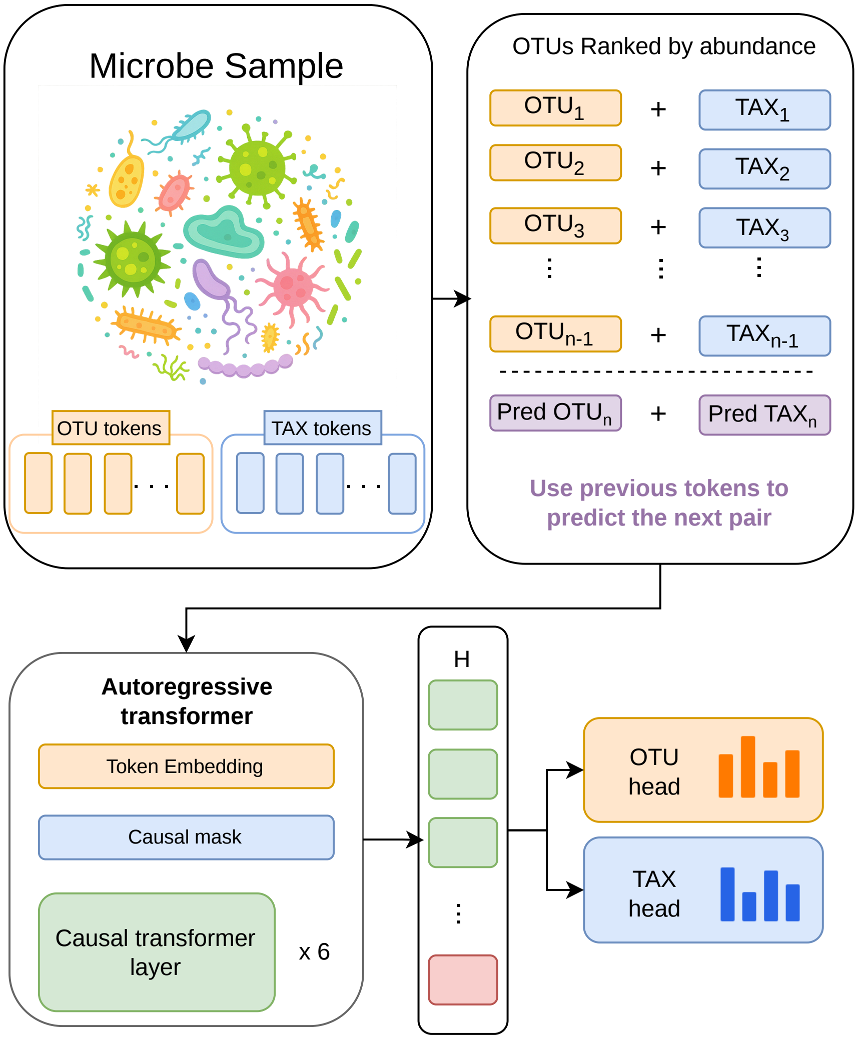


Figure S1: Overview of OTU–Taxa. Each sample is represented as an abundance-ranked sequence of paired OTU and taxonomy tokens. A shared causal Transformer predicts the next OTU and its assigned taxonomy during training. Generation samples the OTU head and supplies the emitted OTU's fixed taxonomy annotation; the learned representation also supports classification and context-based taxonomy-assignment recovery.

where $u = t_i^{(\mathrm{pa}(r))}$ is the assigned parent token during training. Thus,

$$P_\theta\Big(t_i^{(r)} \mid t_i^{(\mathrm{pa}(r))}, \mathbf{x}_{<i}\Big) = \frac{\exp z^{\mathrm{tax}}_{i,t_i^{(r)}}}{\sum_{v \in \mathrm{Child}_r(t_i^{(\mathrm{pa}(r))})} \exp z^{\mathrm{tax}}_{i,v}} \qquad (r \neq K).$$

The extended hierarchy contains one rank-specific unknown token at every rank. For each admissible parent at the preceding rank, the corresponding unknown child is included among the candidates; the unknown tokens also connect to the unknown tokens at deeper ranks. If an annotation becomes unresolved at rank $r$, its target path therefore uses the unknown token at $r$ and at all deeper ranks rather than assigning an incompatible named taxon. This represents the incomplete annotation as a valid path through the extended hierarchy.

With these definitions, the hierarchy-consistent likelihood of the complete taxonomy target at position $i$ is

$$P_\theta(\mathbf{y}_i \mid \mathbf{x}_{<i}) = P_\theta(t_i^{(K)} \mid \mathbf{x}_{<i}) \prod_{r \in \mathcal{R} \setminus \{K\}} P_\theta\Big(t_i^{(r)} \mid t_i^{(\mathrm{pa}(r))}, \mathbf{x}_{<i}\Big).$$

The next-OTU loss is the mean negative log-probability of the next-OTU or EOS targets over the supervised positions:

$$\mathcal{L}_{\mathrm{otu}} = -\frac{1}{|\mathcal{I}_x^{\mathrm{otu}}|} \sum_{i \in \mathcal{I}_x^{\mathrm{otu}}} \log P_\theta(o_i^{\mathrm{lab}} \mid \mathbf{x}_{<i}).$$

The taxonomy-path loss is the negative log-likelihood of the complete target path, summed over ranks and averaged over the same supervised positions:

$$\mathcal{L}_{\mathrm{tax}}^{\mathrm{path}} = -\frac{1}{|\mathcal{I}_x^{\mathrm{tax}}|} \sum_{i \in \mathcal{I}_x^{\mathrm{tax}}} \left[ \log P_\theta(t_i^{(K)} \mid \mathbf{x}_{<i}) + \sum_{r \in \mathcal{R} \setminus \{K\}} \log P_\theta\Big(t_i^{(r)} \mid t_i^{(\mathrm{pa}(r))}, \mathbf{x}_{<i}\Big) \right].$$

The complete pretraining objective is

$$\mathcal{L} = \lambda_{\mathrm{otu}} \mathcal{L}_{\mathrm{otu}} + \lambda_{\mathrm{tax}} \mathcal{L}_{\mathrm{tax}}^{\mathrm{path}}.$$

Here, the nonnegative weights $\lambda_{\mathrm{otu}}$ and $\lambda_{\mathrm{tax}}$ control the relative contributions of the next-OTU and taxonomy-path objectives; the final model used $\lambda_{\mathrm{otu}} = \lambda_{\mathrm{tax}} = 1$.

At inference, the taxonomy head still produces logits over all taxonomy tokens, but predictions are decoded rank by rank. The kingdom prediction is the highest-probability token in $\mathcal{V}_K$. At each lower rank, the admissible set is restricted to the children of the token selected at the preceding rank, and the highest-probability admissible child is selected. This top-down procedure yields one hierarchy-consistent predicted path rather than seven independently selected rank labels.

If the decoder selects the unknown token at rank $r$, that rank and all lower ranks are reported as unresolved, yielding a truncated but valid prediction. Training and decoding therefore apply the same parent–child constraints.

##### M1D. Pretraining Hyperparameters

The final SINTAX-labelled hierarchy-loss checkpoint was trained on the filtered MAP corpus with 1,796,250 training samples, 20,000 validation samples, and 20,000 held-out samples generated from random seed 123. The model had hidden size 256, six Transformer layers, four attention heads, feed-forward size 1024, Gaussian error linear unit (GELU) activations, dropout 0.1, embedding dropout

0.1, layer normalization after summing the input embeddings, a learned abundance-rank embedding, no separate binned-abundance embedding, OTU and taxonomy input embeddings reused as their respective output projections, and maximum sequence length 500. Training used AdamW [3] with learning rate $3 \times 10^{-4}$, weight decay $10^{-3}$, Adam $\epsilon = 10^{-6}$, batch size 256, gradient accumulation over two batches, maximum gradient norm 1.0, and bfloat16 autocast on CUDA. The learning-rate schedule used a 6% linear warmup followed by linear decay. The checkpoint used in downstream analyses was the epoch with the lowest validation loss for the combined OTU next-token loss and hierarchy-aware taxonomy path loss. The final pretraining run used Python 3.12.3, PyTorch 2.7.1, and CUDA 12.8 on one NVIDIA H200 graphics processing unit (GPU). The 24-epoch run completed in 15 h 6 min; its Slurm allocation used eight central processing unit (CPU) cores and 120 gigabytes (GB) of host memory. The model contained 22.57 million trainable parameters.

##### M1E. Abundance-Order Sensitivity

We evaluated the effect of local and tied-rank ordering choices on 1,000 deterministically selected profiles from the final model's held-out test partition. Profiles were restricted to the same maximum of 499 OTUs visible to the model, and five fixed random seeds were used for each perturbation. The exact-tie condition independently permuted OTUs within every block sharing the same read count. The near-tie condition swapped a maximal disjoint set of adjacent OTU pairs when both counts were at most 10 and differed by exactly one read. Each perturbation preserved OTU membership, the OTU–taxonomy pairing, the read-count multiset, and sequence length. Unlike the shuffled-context ablation, these controls preserved sample composition and isolated sensitivity to the induced OTU order.

For each perturbed profile, we compared the mean-pooled non-padding hidden representation used for downstream sample classification with its unperturbed counterpart using cosine similarity. We also compared the final-context next-OTU distributions over the 62,200 real OTUs and the end-of-sample token using Jensen–Shannon divergence. Reported 95% confidence intervals are based on 10,000 profile-level bootstrap resamples after averaging the five seeds within each profile.

As an outcome-level extension, we repeated the community-completion benchmark described in Supplementary Methods M3A on the same 2,000 held-out test profiles. The observed and withheld OTU sets were defined before changing the order, so equal-count OTUs at the split boundary remained on their original side. Within the visible prefix, OTU–taxonomy pairs were independently permuted inside exact-count blocks using five fixed seeds. Counts, sequence length, the order of unequal-count OTUs, candidate OTUs, and withheld targets were unchanged. For each metric, the perturbed-minus-canonical difference was averaged across order seeds within each profile, and 95% confidence intervals were calculated from 10,000 paired profile-level bootstrap resamples.

##### M1F. Matched Taxonomy-Pretraining Controls

We pretrained three conditions from the same seeded initial state: paired OTU–taxonomy inputs with OTU cross-entropy and hierarchy-aware taxonomy-path loss (both weights 1); paired inputs with OTU cross-entropy alone; and OTU-only inputs with OTU cross-entropy alone. SINTAX assignments supplied the taxonomy labels. In the OTU-only condition, the taxonomy-input gate was fixed at zero and taxonomy embeddings remained frozen during both pretraining and downstream fine-tuning. Comparing the first two conditions isolates taxonomy-path supervision with paired inputs fixed; comparing the second and third isolates taxonomy inputs with OTU loss fixed. The first versus third comparison combines these components.

All conditions shared pretraining seed 123, corpus partitions (1,796,250 training, 20,000 validation, and 20,000 test profiles), backbone, abundance ordering, optimizer settings, and profile exposure. Supplementary Methods M1B–M1D describe the shared architecture and training settings. The primary checkpoints were fixed at the end of epoch 24 before downstream outcomes were inspected. All three contained 22,567,680 parameters; in the OTU-only condition, 1,776,384 taxonomy-embedding parameters were frozen, leaving 20,791,296 active parameters. The paired conditions had 22,567,680 active parameters. Training exposure and update counts were matched; active parameter counts and floating-point operation counts therefore need not be identical.

Each epoch used every training profile exactly once. A physical batch size of 256 and accumulation over two microbatches produced 3,508 two-microbatch updates followed by one final update using the remaining 154 profiles. That final accumulation window was normalized by its actual single-microbatch length. Gradients were cleared at epoch boundaries, giving 3,509 updates per epoch and 84,216 updates over 24 epochs in each condition. As a secondary checkpoint-selection check, minimum validation OTU cross-entropy selected epoch 24 in all three conditions; the selected model tensors were identical to the primary checkpoints.

Downstream evaluation used the same 250 tasks from 137 study accessions, with identical fixed outer and group-disjoint inner split sample IDs in all conditions. There were 1,234 outer-fold fits per condition (3,702 overall). Each fit used mean pooling, a 256-unit MLP head, at most 15 epochs, batch size 64, encoder and head learning rates of $5 \times 10^{-5}$ and $5 \times 10^{-4}$, weight decay $10^{-4}$, inverse-frequency class weights, gradient-norm clipping at 1, and BF16 arithmetic. The fine-tuning seed was 42 plus the fold index. Inner-validation balanced accuracy selected the encoder and head with patience 4; both were restored for outer-test evaluation. Supplementary Methods M2B gives the shared classification protocol and metric definitions.

We specified the statistical analysis before examining the results. We first averaged outer-fold scores equally within each task, calculated paired condition differences, averaged task differences within each accession, and finally averaged the 137 accession means equally. Pointwise 95% percentile confidence intervals used 10,000 whole-accession bootstrap draws, shared across conditions and metrics. Two-sided tests used 10,000 shared accession-level sign-flip vectors with the absolute mean paired difference as the statistic. With $b$ sign-flip statistics at least as extreme as observed, the Monte Carlo $p$-value was $(1 + b)/(10{,}000 + 1)$. This test assumes symmetry/exchangeability of the accession-level paired differences around zero. Holm adjustment covered all nine tests jointly (three condition contrasts and three metrics). The analysis seed was 20261102. We also report means with equal weight for each task. The intervals quantify variation across accessions for one shared pretraining seed. Variation between pretraining seeds was not measured.

#### Supplementary Methods M2. Sample Classification

##### M2A. Task Construction and Curation

The MAP-derived sample-classification suite was constructed before model evaluation. Tasks were defined separately within each MAP study accession. For each accession with more than 100 sample records, empty values and the case-insensitive values `not provided` and `na` were omitted during initial metadata enumeration. Candidate fields were required to contain 2–20 observed values, more than 100 eligible annotations, and a smallest-to-largest class-count ratio of at least 0.10.

All model inputs used the common filtering and abundance-order preprocessing described in Supplementary Methods M1A.

Fields that passed these filters were curated in two sequential per-study requests through OpenRouter using the `meta-llama/llama-3-70b-instruct` model and temperature 0.2. Each request contained the metadata-field names and their complete class-value count dictionaries. The first pass selected biologically or environmentally meaningful classification labels using a maximum of 1,024 output tokens. The second pass used a maximum of 1,200 output tokens and targeted constant, missing-dominated, protocol-related, and redundant fields. Responses were parsed into field-to-class-count dictionaries.

Retained study–field pairs were joined to samples from the corresponding MAP accession that had filtered OTU profiles. During final label export, surrounding whitespace was removed and the case-insensitive values `na`, `n/a`, `nan`, `none`, `null`, `unknown`, `missing`, and `not provided` were omitted. Binary tasks required exactly two observed classes with at least 100 samples per class, whereas multiclass tasks required more than two classes with at least 50 samples per class. The resulting 639 task identities, study accessions, metadata fields, sample counts, and class counts were frozen in the released benchmark manifest, available in the OTU–Taxa paper-artifacts dataset (`https://huggingface.co/datasets/bio-ontology-research-group/otu-taxa-paper-artifacts`). To support reproducibility of LLM-assisted metadata curation, the selection criteria described above, exact prompt templates, recorded model settings, and the saved intermediate curation dictionaries are provided in the OTU–Taxa source repository (`https://github.com/bio-ontology-research-group/otu-taxa-autoregressive`).

As an abbreviated first-pass example, the request for study ERP006059 used the system instruction "You are an expert in biological data curation." The user prompt instructed the model: "For each field, decide if it is a meaningful candidate for a biological or environmental classification label." The metadata supplied included the following entry:

```
{
  "sample_body_site": {
    "UBERON:feces": 534,
    "UBERON:saliva": 286
  }
}
```

The body-site field was retained in the saved curation outputs. Per-sample labels were subsequently obtained from the corresponding MAP metadata; after matching samples to available OTU profiles and applying the final label filters, this task contained 512 feces and 285 saliva samples. The full per-study request included other eligible metadata fields. The complete prompt templates for both curation passes can be inspected at `https://github.com/bio-ontology-research-group/otu-taxa-autoregressive/blob/127c5f15e0f49f1c8aba98dd3cdf5f44a2dc0193/configs/classification/task_curation_prompts.json`; the saved curation outputs are available in the source repository linked above.

The 639-task panel comprised 449 binary and 190 multiclass tasks. The median task contained 593 samples (range 208–6,837), the median minimum class count was 176 (range 50–2,014), and tasks contained 2–18 classes.

The classification benchmark evaluates in-domain prediction of withheld metadata labels using profiles drawn from the MAP corpus used for self-supervised pretraining. Consistent with this setting, the overlap audit found substantial exposure across biological identifiers. Among profiles with available identifiers, BioSample overlap with the pretraining training partition was 98.20% and 97.89%, and study-qualified subject overlap was 99.51% and 99.72%, for the 639-task and 250-task panels, respectively. Identical model inputs under different profile IDs occurred in 347 and 72 cases. Among evaluation profiles whose exact IDs were absent from that training partition, 179/6,712 and 49/2,380 had training-profile OTU-set Jaccard similarity $\geq 0.90$.

The benchmark's shared MAP OTU reference can be used to compare methods with different training histories, including methods pretrained on other corpora. Each evaluation study can be used separately through its associated OTU profiles and metadata labels. Additional compatible datasets can also be added by mapping their sequences to the MAP OTU reference, providing a common feature vocabulary for comparison. The term “in-domain” describes the relationship between our pretrained models and the evaluated profiles; it does not restrict the benchmark to models trained on MAP.

##### M2B. Fine-Tuning and Baseline Protocols

Cross-validation partitions were constructed at the level of subjects or repeated experimental units whenever reliable identifiers were available. These identifiers represented subjects, patients, donors, animals, or experimental units such as plots, tanks, and pens. We refer to these as disjoint folds: all samples belonging to the same subject or repeated experimental unit were assigned to the same fold, preventing related observations from appearing in both the training and test partitions. The identifiers were used only to construct the folds and were never included as predictive features.

Among the final 639 tasks, 380 lacked a reliable subject or repeated-unit identifier, three could not form a valid three- or five-fold disjoint partition, and 256 supported valid disjoint folds. A pre-fitting coverage check found complete required-sample representation across OTU–Taxa, Pope-MAP, the Microbial General Model (MGM), and Waypoint for 250 of these 256 tasks. In the other six tasks, some samples contained no genus-resolved OTU after conversion to MGM’s abundance-ranked genus representation. Waypoint introduced no additional missing profiles. These tasks were excluded from the matched primary comparison rather than reducing their folds. This coverage rule was applied before model fitting and did not depend on downstream performance.

The primary 250 tasks comprise 242 tasks with five disjoint outer folds and eight with three disjoint outer folds, used when the number or class distribution of independent units did not support five valid folds. Hyperparameter selection and early stopping followed the same disjointness constraint in inner validation partitions drawn exclusively from the outer-training data, and all methods used the same recorded outer and inner partitions. The remaining 389 tasks used stratified sample-level folds and are reported as a supplementary comparison. The released task manifest records the study accession, class-label field, grouping field and type, and sample-to-fold assignments.

Table S1 records the pretraining and transfer configuration of each pretrained comparator. All were adapted from the same filtered MAP source collection, but retained their model-specific token definition, architecture, pretraining objective, sequence length, and pooling rule. Representation-specific filtering accounts for the different numbers of profiles retained for MGM and Waypoint.

Table S1: Configuration of the pretrained microbiome models used for the MAP comparison. Profile counts are train/validation/test after model-specific conversion. Pope-MAP retains Pope’s abundance-ranked representation and generator–discriminator objective, with the MAP-specific optimization described below.

| Comparison axis | OTU–Taxa | Pope-MAP | MGM | Waypoint |
|---|---|---|---|---|
| MAP profiles | 1,796,250 / 20,000 / 20,000 | 1,796,250 / 20,000 / 20,000 | 1,713,830 / 19,045 / 19,043 | 1,602,693 / 17,825 / 17,769 |

Continued on next page

Table S1 continued

| Comparison axis | OTU–Taxa | Pope-MAP | MGM | Waypoint |
|---|---|---|---|---|
| Vocabulary and representation | 62,200 97%-identity OTUs paired with 6,929 taxonomy nodes; raw-count abundance order | 62,200 OTUs initialized with 100-dimensional Global Vectors for Word Representation (GloVe) vectors; raw-count abundance order | 2,567 genus tokens; OTU abundances summed by genus, normalized, and ranked | 3,739 taxonomy tokens; genus with higher-rank fallback, ordered by token-specific abundance $z$-score |
| Maximum input | 500 OTU–taxonomy pairs | 512 OTUs plus a classification (CLS) token | 512 positions including special tokens | 512 positions including special tokens |
| Architecture and parameters | Six-layer autoregressive Transformer; hidden size 256; four heads; 22.57M | Efficiently Learning an Encoder that Classifies Token Replacements Accurately (ELECTRA) generator: four layers, four heads, hidden size 200, 13.88M; discriminator: five layers, five heads, hidden size 200, 14.16M | Eight-layer Generative Pre-trained Transformer 2 (GPT-2); hidden size 256; eight heads; 7.11M | Waypoint-6m GPT-2; eight layers; hidden size 256; four heads; 7.41M |
| Pretraining objective | Causal next-OTU prediction plus hierarchy-aware taxonomy-path prediction; losses weighted 1:1 | Masked-OTU generation followed by replaced-token discrimination | Causal next-genus-token prediction | Causal next-taxonomy-token prediction |
| Pretraining optimization | AdamW; learning rate $3 \times 10^{-4}$; effective batch 512; 24 epochs | AdamW; generator learning rate $5 \times 10^{-5}$, effective batch 400, 10 epochs; discriminator learning rate $2 \times 10^{-5}$, batch 400 | AdamW; learning rate $10^{-3}$; batch 256; 9.19 epochs | AdamW; learning rate $10^{-3}$; batch 256; 27 epochs |

Continued on next page

Table S1 continued

| Comparison axis | OTU–Taxa | Pope-MAP | MGM | Waypoint |
|---|---|---|---|---|
| Checkpoint selection | Lowest combined validation objective | Generator epoch 10; discriminator step 600, selected by the lowest fixed-validation balanced loss | Lowest validation loss | Lowest validation loss after 43.27M profile presentations |
| Downstream readout | Mean pooling and 256-unit multilayer perceptron (MLP) head | CLS pooling and 200-unit MLP head | EOS pooling and 256-unit MLP head | EOS pooling and linear head |
| Downstream fitting | Identical saved outer and inner folds; full encoder fine-tuning with AdamW; encoder/head learning rates $5 \times 10^{-5}/5 \times 10^{-4}$; weight decay $10^{-4}$; batch size 64; at most 15 epochs; class-weighted cross-entropy; gradient clipping at 1.0; inner-validation balanced-accuracy selection with patience four. Neural hyperparameters were fixed before evaluation rather than searched per task. | | | |

To construct the Pope-MAP comparator, we first followed the published implementation as closely as possible: samples were represented by up to 512 abundance-ranked OTUs, initialized with 100-dimensional GloVe embeddings, and processed by a ten-layer, ten-head masked-OTU generator followed by a five-layer, five-head replaced-token discriminator. When transferred to the substantially larger MAP corpus, this configuration did not show a sustained reduction in the held-out pretraining objective.

We therefore evaluated controlled modifications while retaining Pope's abundance-ranked representation and ELECTRA-style pretraining design. For the generator, we compared trainable GloVe initialization, trainable random initialization, delayed unfreezing of GloVe embeddings, and a reduced embedding learning rate. The selected configuration used a smaller four-layer, four-head generator with GloVe embeddings trainable from the beginning and was trained for ten epochs. For the discriminator, we evaluated different generator-checkpoint schedules, frozen or slowly updated OTU embeddings, unweighted or class-balanced binary cross-entropy, and different training exposures. The selected discriminator used the fixed epoch-10 generator from its first step, assigned equal weight to replaced and original tokens, updated the real-OTU embeddings at one tenth of the Transformer learning rate, and selected step 600 using held-out balanced loss.

We refer to the resulting model as Pope-MAP because it preserves Pope's representation, generator–discriminator objective, and downstream CLS readout, but uses the most stable MAP-trained implementation identified in these experiments rather than reproducing the published checkpoint. Additional model-specific optimization could potentially improve any of the pretrained comparators. Our objective was therefore to preserve their published representation and learning strategy where possible, train them on the same MAP source collection, and evaluate them using the same downstream folds and fine-tuning budget.

Using the recorded partitions, neural sample classification reloaded the pretrained encoder for each outer fold and used the approximately 15% inner validation partition for checkpoint selection. The held-out outer fold remained untouched until evaluation. Each model retained the pooling rule and classification head shown in Table S1. The OTU–Taxa and MGM heads were LayerNorm–Linear(256)–GELU–Dropout(0.1)–Linear($C$), where $C$ is the number of task classes; Pope-MAP retained its 200-unit MLP and Waypoint retained its released linear head. The common full-fine-tuning protocol used the settings in Table S1, without warm-up or a learning-rate scheduler. Class weights were estimated only from the corresponding inner-training partition.

Frozen-head and frozen-probe variants were also evaluated as adaptation controls using the same partitions and fold-local preprocessing; the reported OTU–Taxa classification results use full fine-tuning.

On the same outer and inner partitions, and following established microbiome-classification practice [4, 5], Logistic Regression and Random Forest were trained on sample-by-OTU relative abundances. Logistic Regression used the `saga` solver, an $\ell_2$ penalty, `class_weight="balanced"`, and $C \in \{0.01, 0.1, 1, 10, 100\}$. Random Forest used balanced class weights. Its 24-setting search crossed 300 or 600 trees, square-root or $\log_2$ feature sampling, minimum leaf sizes of 1, 3, or 5, and either unrestricted depth or a maximum depth of 20. Hyperparameters were selected on the corresponding inner validation partition. The selected model was then refitted on the complete outer-training partition and evaluated once on the untouched outer fold.

Supplementary Table S6 reports separate five-fold subject-disjoint reruns for American Gut Project (AGP) inflammatory bowel disease (IBD) and Human Microbiome Project (HMP) body site. The classical models use relative-abundance OTU features and the full hyperparameter search described above, with subject-disjoint inner validation partitions drawn from each outer-training fold.

##### M2C. Critical-Difference Ranking

Across-task method rankings use the common-task subset. For each metric, ranks are computed from task-level mean cross-validation scores, with rank 1 assigned to the highest-scoring method and ties assigned average ranks. A Friedman omnibus test was applied separately to balanced accuracy, area under the receiver operating characteristic curve (AUROC), and area under the precision–recall curve (AUPR), and the three resulting $p$-values were adjusted by the Holm procedure. Nemenyi critical differences (CDs) were then computed at $\alpha = 0.05$ [6] as $\text{CD} = q_\alpha \sqrt{k(k+1)/(6N)}$, where $k$ is the number of methods and $N$ is the number of paired tasks for the corresponding metric.

The statistical unit was the prediction task. Each study–metadata-field combination had a distinct label definition and independently fitted cross-validation analysis and was therefore treated as a separate task. The 250 primary tasks spanned 137 study accessions; 63 accessions contributed more than one task, and these tasks may share samples and study-level characteristics. We therefore performed a sensitivity analysis in which task ranks were first averaged within accession before the Friedman tests. For both task- and accession-level analyses, a Nemenyi connection denotes failure to detect the specified rank difference and should not be interpreted as evidence that two methods are equivalent.

The primary analysis uses the 250 tasks evaluated with disjoint folds. Supplementary Results R3 report the 389 stratified sample-level tasks separately.

#### Supplementary Methods M3. Sample Generation and Disease-Ecology Readouts

##### M3A. Generation and Community-Completion Baselines

For unconditional generation, the final hierarchy-aware checkpoint was sampled with ten fixed seeds, producing 3,000 profiles per seed. Each replicate was evaluated against the same 5,000 held-out MAP profiles. The global empirical retrieval reference randomly samples profiles from the training partition, whereas the same-study reference samples a training profile from the study of the paired held-out target. Runs from the same BioSample were excluded. A same-study training profile was available for 2,904 of 3,000 targets (96.8%); no environment fallback was used. We report the mean and sample standard deviation across the ten stochastic generation or retrieval replicates. Exact-copy and nearest-profile comparisons use the complete 1,796,250-profile training partition.

For each replicate, richness is the number of OTUs in a profile. Richness Jensen–Shannon (JS) divergence compares the generated or retrieved and held-out richness histograms using 40 evenly spaced bin edges from zero to one above the largest observed richness; divergence is calculated with natural logarithms. OTU prevalence is the fraction of profiles containing each OTU, and prevalence Pearson correlation is calculated across OTUs observed at least once in the held-out reference. Co-occurrence metrics use the 200 most prevalent OTUs in the held-out reference. Binary profile-by-OTU presence matrices are converted to pairwise OTU Pearson-correlation matrices, and their upper triangles are compared between each generated or retrieved set and the held-out reference by Pearson correlation and root mean squared error (RMSE). A generated profile is counted as an exact copy when its set of OTUs is identical to any profile in the training partition, equivalently when its maximum training-profile Jaccard similarity is one.

The conditional community-completion benchmark compares all methods on the same held-out test samples, observed prefixes, target OTUs, and candidate OTU sets. After deterministic shuffling with seed 42, we evaluated the first 2,000 test profiles containing at least eight OTUs. Profiles exceeding the model context were restricted to their 499 most-abundant OTUs, together with the beginning-of-sample token, before the retained profile was divided into observed and withheld halves. This restriction affected 104 of the 2,000 evaluated profiles. Observed OTUs are removed from the candidate set before ranking. For OTU–Taxa, every remaining candidate is scored from the single next-token distribution conditioned on the fixed observed half-profile. The prefix is not extended with predicted tokens, so recall and ranking over the withheld half quantify one-step conditional association rather than sequential reconstruction of the suffix. The evaluation pool was the 20,000-sample test partition, disjoint from the validation partition used for checkpoint selection.

The global-popularity baseline ranks candidate OTUs by their unigram frequency in the MAP training corpus. For the word2vec co-occurrence baseline [7], each MAP sample is an OTU sentence ordered by abundance; samples longer than 256 OTUs are truncated to the most abundant OTUs. We trained 100-dimensional skip-gram embeddings with negative sampling using five negatives per positive pair, where the context OTU for a center OTU is sampled from the same microbiome sample and negatives are drawn from the unigram distribution raised to the 0.75 power. The word2vec score for candidate OTU $o$ is the summed cosine similarity between the candidate embedding and the embeddings of the observed-prefix OTUs: $s_{\mathrm{w2v}}(o; \mathcal{O}_{\mathrm{obs}}) = \tilde{e}_o^\top \sum_{o' \in \mathcal{O}_{\mathrm{obs}}} \tilde{e}_{o'}$, where $\tilde{e}$ denotes a unit-length word2vec embedding and $\mathcal{O}_{\mathrm{obs}}$ is the observed-prefix OTU set. The learned-embedding co-occurrence baseline uses the same scoring rule, replacing the word2vec embeddings with the normalized OTU embedding table learned by OTU–Taxa during pretraining.

The sample-retrieval baseline identifies the training profile with the greatest exact Jaccard similarity to the observed test prefix. All training runs sharing the query's BioSample accession are excluded. Unobserved OTUs in the retrieved profile are ranked by their abundance order, followed by remaining candidates in training-set popularity order. Equal-Jaccard neighbours are resolved by the fixed training index. This baseline uses only the observed prefix and the same 62,200 candidate OTUs used by OTU–Taxa.

For a cutoff $k$, Recall@$k$ is the fraction of withheld OTUs recovered among the top $k$ candidates, and normalized discounted cumulative gain at $k$ (NDCG@$k$) rewards correct candidates more when they occur earlier in the ranked list. Area under the precision–recall curve (AUPRC) summarizes the complete candidate ranking. We use "autoregressive (AR) completion" for the OTU–Taxa next-token ranking conditioned on the observed prefix.

For each completion metric, we report the sample mean and a 95% percentile-bootstrap confidence interval from 10,000 sample-level resamples. The paired comparison with sample retrieval resamples the same profiles and summarizes the per-profile difference between OTU–Taxa and retrieval. These intervals quantify evaluation-sample uncertainty for one fixed trained checkpoint rather than variation across training runs.

**Sequential prediction comparison.** We additionally compared four prediction procedures using the same checkpoint and 2,000 held-out profiles: fixed-prefix ranking, sequential greedy prediction, sequential sampling, and frozen sampling. Sequential procedures append each predicted OTU and its fixed taxonomy annotation before predicting again. Frozen sampling draws without replacement from the initial prefix distribution. Every procedure outputs exactly the number of withheld OTUs, excluding observed and previously selected OTUs; early stopping is disabled. Recovery is measured using recall and Jaccard similarity. Sampling uses temperature 1 and three seeds, averaged within each profile. Confidence intervals use 10,000 paired profile-bootstrap resamples. The existing retrieval and popularity baselines use the same output budget.

##### M3B. BugSigDB Reference Construction

The external IBD-related literature reference was constructed from the full BugSigDB comma-separated-value (CSV) export [8] downloaded on 30 May 2026 at 13:25 Coordinated Universal Time (UTC), as recorded in the export header. We retained signature rows whose host species was *Homo sapiens*, whose condition contained "inflammatory bowel disease", "Crohn", or "ulcerative colitis" case-insensitively, and whose reported abundance in the annotated Group 1 contrast was either increased or decreased. No additional restriction was placed on study design, body site, sequencing method, or the definitions of Group 0 and Group 1; the retained set therefore includes both disease-versus-control and within-IBD clinical contrasts.

Taxa were read from the export's MetaPhlAn lineage field and mapped to genus by extracting the `g__` label. Species- and strain-level entries therefore contributed at their annotated genus, whereas entries without a genus label were excluded. A genus was counted at most once within a signature after this collapse. Distinct signature rows were retained even when they originated from the same study, so the reference weights signatures rather than studies and does not apply study-level deduplication or reweighting. These rules retained 186 directional signatures from 33 studies, comprising 157 genera with at least one increased report and 139 genera with at least one decreased report; a genus could occur in both sets. The pooled score $\mathrm{net}_{\mathrm{IBD}}(g)$ is the number of retained signatures reporting genus $g$ as increased in Group 1 minus the number reporting it as decreased. Genera with a zero pooled score were omitted from direction-based comparisons.

##### M3C. Within-Cohort Disease-Ecology Readouts

**Cohorts and named comparisons.** All within-cohort disease-ecology analyses used profiles represented in MAP, with task definitions fixed before model scoring. The two highlighted examples were the watering-regime contrast from accession SRP098753 (318 control and 298 drought profiles) and the environment-medium contrast from accession SRP250131 (656 lake-water and 176 stream-water profiles). For the literature-facing IBD comparison, the American Gut Project label table [9] contained 18,585 profiles (17,861 controls and 724 IBD cases), and the Halfvarson cohort (ERP020401) [10] contained 566 profiles (57 controls and 509 IBD cases). Model and count readouts used the same cohort definitions; the model-score subsampling and count-analysis coverage are specified below, and the external genus directions were obtained from the BugSigDB construction in Supplementary Methods M3B.

We evaluated 100 prespecified binary contrasts, each defined by one selected metadata field within a study accession. The two selected values of that field divided samples into groups denoted neg and pos to orient the comparison. For the count-based analysis, OTU relative abundances were summed by genus, and genera observed in at least 10 samples were tested. The observed log fold change (logFC) was the natural logarithm of the ratio between the pos- and neg-group mean relative abundances, with $10^{-9}$ added to both means. Validated subject or repeated-unit identifiers were available for 43 contrasts. Labels from mixed-label units were permuted within unit, whereas class-constant units were reassigned as whole units; nine partially crossed contrasts combined these components. The remaining 57 contrasts had no validated repeated-unit identifier and used sample-level permutations preserving the observed class sizes. Four of the 43 mapped contrasts contained only singleton units, for which unit- and sample-level permutations are equivalent. Each contrast used 10,000 permutations. Two-sided permutation $p$-values compared the absolute permuted and observed effects, and Benjamini–Hochberg correction was applied separately within each contrast across its tested genera to control the false-discovery rate (FDR). The two adjustments address different hypothesis families: Holm controls family-wise error across the three global classification tests, whereas Benjamini–Hochberg controls the false-discovery rate among the genera tested separately within each contrast.

To bound autoregressive inference cost, model-derived scores were estimated from a seed-42 selection of at most 400 profiles per group. Each profile was evaluated over teacher-forced prefixes of up to 300 OTUs, and the resulting next-OTU probabilities were summed by genus without masking OTUs already present in the prefix. Count-based abundance estimates used all eligible profiles. For group $c$, the heuristic model abundance-rank score uses

$$R_c(g) = \frac{\sum_{x \in c} \sum_i i P_\theta(g \mid \mathbf{x}_{<i})}{\sum_{x \in c} \sum_i P_\theta(g \mid \mathbf{x}_{<i})}, \qquad S(g) = R_{\text{neg}}(g) - R_{\text{pos}}(g).$$

Positive values indicate that the model concentrates more genus probability at earlier abundance-ranked positions in the pos group.

Model–count agreement was calculated for common genera with count-side $q \leq 0.05$ and a nonzero abundance difference. AUROC uses the direction of the observed abundance difference as the binary genus label and $S(g)$ as the predictor, while Spearman correlation compares their signed values. Of the 100 contrasts, 98 produced genus-mapped model scores and 78 contained at least 10 selected genera with a defined AUROC. The top-half and top-quarter analyses formed nested subsets using each genus's pooled observed mean relative abundance, weighted by the pos- and neg-group sample counts and ranked among all genera shared by the count and model tables for that contrast. The subsets retained selected genera at or above the 50th and 75th percentiles, respectively. AUROC is undefined when a subset contains only one abundance-change direction; this did not occur at any of the three thresholds among the 78 retained contrasts.

Because genus selection and abundance-change direction were derived from the same count contrast, these AUROCs summarize model–count agreement conditional on the selected genera rather than agreement across all observed genera. To test whether the mean conditional AUROC exceeded chance, the selected genus sets and count-derived directions were held fixed while model-score ranks were permuted among genera within each contrast. Each of 10,000 permutation replicates was summarized using the equal-contrast mean AUROC. Confidence intervals were obtained from 10,000 study-accession bootstrap resamples, retaining all eligible contrasts from each sampled accession. The 78 contrasts represented 74 study accessions, including four accessions contributing two contrasts.

#### Supplementary Methods M4. Context-Based Taxonomy-Assignment Recovery

##### M4A. Sequence-Degradation Benchmark

The sequence-degradation benchmark measures context-based recovery of classifier assignments as sequence-local evidence is reduced. The clean representative OTU sequences define the full-sequence SINTAX and Quantitative Insights Into Microbial Ecology 2 (QIIME2) reference assignments. Degraded versions of the same sequences are classified again and scored in the corresponding reference space. SINTAX and RAxTax are evaluated against the SINTAX assignments, whereas QIIME2 is evaluated separately against the QIIME2 assignments.

**Purpose of the experiment.** Sequence-based taxonomy classifiers assign labels using the query sequence and annotated references. Community context also provides information about the taxonomy expected at a position in an abundance-ranked profile. OTU–Taxa learns these relationships within a fixed MAP OTU vocabulary. We keep the observed community context unchanged while progressively degrading the target representative sequence, and measure recovery of its intact-sequence classifier assignment. The aim is to assess community context as an additional source of taxonomic information for reference-mapped communities, particularly at higher ranks. The context-only readout uses preceding community members, with the target sequence, OTU identity, and taxonomy withheld at inference.

Eligible OTUs were required to have a resolved genus- or species-level assignment in the taxonomy table for the corresponding label source. For the SINTAX-aligned context comparison, 5,666 OTUs had both eligible full-sequence SINTAX labels and held-out MAP samples containing them for context-only evaluation. These OTUs produced 43,215 sample-specific OTU–Taxa predictions before OTU-level majority aggregation. For the QIIME2-labeled analysis, the matched context-only evaluation included 4,168 OTUs and 31,627 sample-specific predictions. RAxTax was rescored on the same 5,666 OTUs and against the same full-sequence SINTAX assignments used for the SINTAX-aligned comparison.

The SINTAX checkpoint used the configuration in Supplementary Methods M1D. The matched QIIME2 checkpoint used the identical MAP split, OTU vocabulary, maximum length, architecture, optimization schedule, hierarchy-aware taxonomy-path objective, equal loss weights, random seed 123, 24 epochs, and batch size 256. Only its QIIME2-derived taxonomy hierarchy and prediction head differed: 7,699 observed nodes, 7,705 nodes after adding rank-specific unknowns, and 7,709 taxonomy tokens after special tokens.

The degradation grid is summarized in Table S2. Each substitution or deletion condition uses one deterministic replicate per OTU. For substitutions, each canonical base position $\{A, C, G, T\}$ is independently selected with probability $p$; selected bases are replaced by one of the other three canonical bases. Non-canonical bases are left unchanged. For deletions, each position is independently removed with probability $p$. For truncation, the central fragment of the requested length is retained; if a sequence is shorter than the requested fragment length, the complete sequence is retained. Random number generators are seeded by the global seed and the degradation condition, making the sequence files deterministic.

SINTAX predictions [11] were generated with VSEARCH v2.21.1 [12], using the SINTAX-formatted SILVA v123 16S reference [13] downloaded from the USEARCH SINTAX reference collection (`https://www.drive5.com/usearch/manual/sintax_downloads.html`) and a confidence cutoff of 0.8. QIIME2 labels were generated with QIIME2 v2026.1.0 and `q2-feature-classifier` v2026.1.0 [14, 15]. The classifier was trained with `fit-classifier-naive-bayes` on a separately prepared, full-length SILVA v123-derived 16S reference from which 2,058 species had been

Table S2: Controlled sequence-degradation grid used for the context-based taxonomy-assignment recovery benchmark.

| Classifier | Family | Levels | Sequence effect |
|---|---|---|---|
| SINTAX | Substitution/deletion | 10, 12, 14, 16, 18, and 20% | Base identity perturbed or positions removed |
| SINTAX | Centered truncation | 25, 30, 50, 100, 150, 200, 250, 300, and 500 bp | Central fragment retained |
| QIIME2 | Substitution/deletion | 1, 3, 5, 10, 12, 14, 15, 16, 18, and 20% | Base identity perturbed or positions removed |
| QIIME2 | Centered truncation | 25, 30, 50, 75, 100, and 150 bp | Central fragment retained |
| RAxTax | Substitution/deletion | 10, 12, 14, 16, 18, 20, 25, 30, and 35% | Base identity perturbed or positions removed |
| RAxTax | Centered truncation | 25, 30, 50, 100, 150, 200, 250, 300, and 500 bp | Central fragment retained |

withheld. The reference contained 1,613,243 sequences (median length 1,384 nt) and was not restricted by in silico primer extraction. Predictions used `classify-sklearn`, a confidence threshold of 0.7, and automatic read-orientation detection. These labels were also used to train the secondary QIIME2-labeled OTU–Taxa checkpoint, and the QIIME2 degradation curves are compared with this QIIME2-labeled context-only readout. RAxTax [16] is included as a sequence-classifier degradation control. Its database was built from the SILVA 16S reference sequences used for the sequence-degradation benchmark, allowing it to serve as an additional sequence-classifier robustness reference under controlled perturbations. Its predictions were evaluated on the SINTAX-aligned 5,666-OTU set against the original full-sequence SINTAX assignments and compared with the SINTAX-trained context-only OTU–Taxa score. All classifier outputs are parsed to the common rank order $k, p, c, o, f, g, s$ and evaluated with per-rank accuracy and hierarchical rank accuracy (rankACC) relative to these classifier-derived assignments. Let $\mathcal{R}_i$ be the ordered set of ranks with available reference annotations for OTU $i$, from kingdom toward species, and let $d_i = |\mathcal{R}_i|$. Missing reference annotations are excluded from scoring, while available finer-rank annotations remain evaluable. Let $m_i$ be the number of consecutive correct predictions along this ordered set before the first disagreement. Then

$$\text{rankACC}_i = \frac{m_i}{d_i}.$$

A missing prediction at an evaluable rank counts as a disagreement. Per-rank agreement uses OTUs with an available reference annotation at that rank; complete-path agreement requires agreement at every available reference rank. Mean rankACC ranges from 0 to 1 and gives partial credit to paths that remain correct through higher ranks before diverging at a finer rank.

The QIIME2 sequence-truncation analysis therefore measures how this specific full-length-trained classifier responds when progressively shorter query fragments are supplied. Training a classifier on primer-extracted sequences from the sequenced region can change taxonomic accuracy [15], as can the reference composition, confidence threshold, and read-orientation setting. Accordingly, the SINTAX and QIIME2 curves locate degradation within each fixed workflow relative to its own clean assignment; their side-by-side presentation is not an absolute comparison of the two classifiers.

For context-only OTU–Taxa prediction, evaluation uses held-out test samples in which the selected OTU is observed. At each evaluated occurrence, the model sees the abundance-ordered prefix before the target position; the target OTU identity and target taxonomy are not provided at the prediction step. Up to ten held-out occurrences are evaluated per selected OTU, and fewer are used when an OTU has fewer available held-out occurrences. Occurrence-level predictions are decoded as hierarchy-consistent paths and then aggregated to one OTU-level label at each rank by rank-specific majority vote, matching the one-assignment-per-OTU structure of sequence-classifier outputs. Because votes are taken separately at each rank, the aggregate labels are not required to reproduce one complete occurrence-level decoded path; rankACC is scored as consecutive agreement from the root.

The context ablations were implemented as follows. Full context retained both observed channels. OTU-only context replaced taxonomy tokens after the beginning-of-sample token with padding embeddings, and taxonomy-only context did the converse. Shuffled context retained the target and its reference label but replaced the prefix with one from another held-out sample of similar prefix length, preferentially excluding prefixes that already contained the target OTU. The global-prior control predicted one rank-conditional path obtained by following the most frequent child at each rank in the pretraining partition. The visible-context analysis retained the first 10%, 25%, 50%, 75%, or 100% of real prefix positions after the beginning-of-sample token, with at least one position retained when a nonempty prefix was available.

### Supplementary Results

#### Supplementary Results R1. Pretraining Optimization

Figure S2 reports the saved joint and component losses over 24 pretraining epochs.

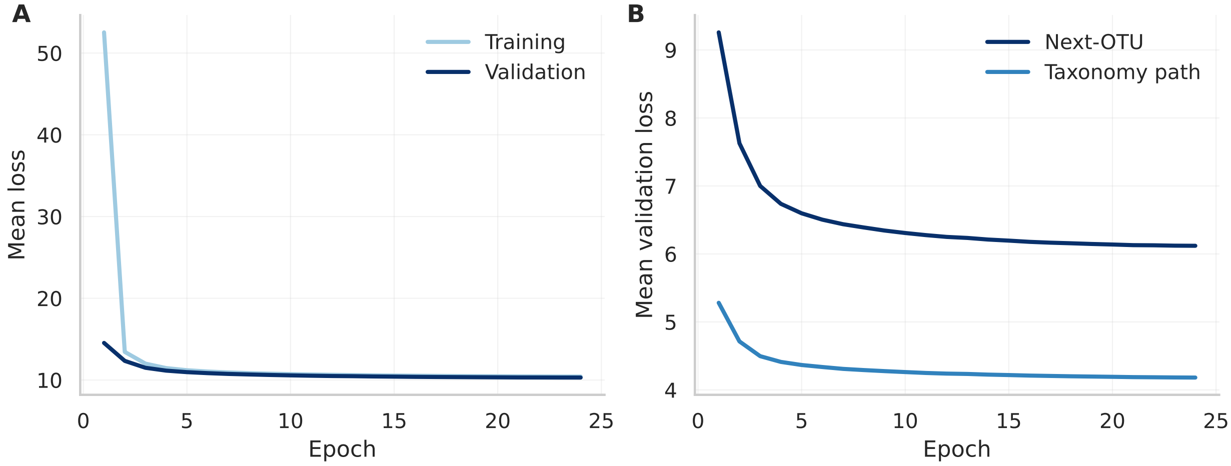


Figure S2: Pretraining loss for the final hierarchy-aware OTU–Taxa model. **A**, mean training and validation joint loss over 24 epochs. **B**, mean validation next-OTU and taxonomy-path loss components. Values are the saved epoch means.

#### Supplementary Results R2. Abundance-Order Sensitivity

Production sorting was stable: equal-count OTUs retained their order in the corresponding MAP source record rather than being sorted by vocabulary index or randomized during tokenization. Among adjacent OTUs within exact-count blocks, vocabulary indices increased in 50.2% of pairs and decreased in 49.8%, providing an empirical check that tied OTUs were not ordered monotonically by vocabulary index.

In the 1,000 held-out profiles, 97.1% contained at least one exact count tie, and 75.3% of model-visible OTUs belonged to an exact-tie block. Ties were concentrated beyond the high-abundance head: the fraction of tokens in exact ties was 5.1% at ranks 1–10, 31.0% at ranks 11–30, 75.2% at ranks 31–100, and 97.5% at ranks 101–499. OTUs with counts of at most 10 comprised 42.8% of the visible tokens.

The mean-pooled sample representation changed little under either order perturbation (Table S3). Local adjacent one-read swaps also produced only a small change in the final-context next-token distribution. By contrast, permuting every exact-count block was a broader intervention that moved nearly half of the visible tokens and produced a larger change in the autoregressive distribution. Sample embeddings therefore changed little under either perturbation, while next-token predictions were less sensitive to local low-count swaps than to extensive reordering of the tied tail. These controls measure sensitivity to induced order, not rank reproducibility under repeated sequencing or rarefaction.

Because the full exact-tie perturbation changed the next-token distribution, we next examined whether this sensitivity altered community completion. Among the 2,000 completion queries, 1,675 (83.8%) contained at least one exact-count tie in the visible prefix, and 54.6% of visible OTUs belonged to tied blocks. The alternative orderings moved 25.0% of visible OTUs on average while preserving the observed and withheld community members.

Community-completion performance did not decrease under the alternative tie orders (Table S3). Mean paired changes ranged from +0.0033 for Recall@10 to +0.0363 for NDCG@100, and the standard deviation across the five order-seed means was at most 0.0016. Sample retrieval remained stronger for Recall@10 and NDCG@10/50, whereas OTU–Taxa retained higher Recall@50/100. Under the alternative orders, the detailed NDCG@100 and AUPRC comparisons shifted in favor of OTU–Taxa. The secondary tie order therefore affected precise ranked predictions, but the aggregate early-ranking-versus-broader-recovery pattern was preserved.

Table S3: Sensitivity of the final model to abundance-order perturbations. **A**, representation and next-token sensitivity on 1,000 held-out test profiles; values are profile means with 95% profile-bootstrap confidence intervals after averaging five fixed perturbation seeds. Jensen–Shannon divergence is reported in natural-log units. **B**, community completion on 2,000 held-out test profiles. Canonical and tie-permuted values are means with 95% profile-bootstrap confidence intervals; the five tie orders were averaged within each profile before resampling. Changes are perturbed-minus-canonical paired means with 95% profile-bootstrap confidence intervals.

**A. Representation and next-token sensitivity**

| Perturbation | Tokens moved | Embedding cosine | Next-token JS |
|---|---|---|---|
| Exact-count blocks | 47.4% [46.0, 48.8] | 0.99871 [0.99855, 0.99885] | 0.286 [0.272, 0.301] |
| Adjacent low-count one-read pairs | 10.7% [10.3, 11.2] | 0.999976 [0.999969, 0.999981] | 0.0102 [0.0091, 0.0115] |

**B. Community-completion outcome sensitivity**

| Metric | Canonical [95% CI] | Tie-permuted [95% CI] | Change [95% CI] |
|---|---|---|---|
| Recall@10 | 0.1005 [0.0954, 0.1057] | 0.1038 [0.0987, 0.1090] | +0.0033 [0.0027, 0.0039] |
| Recall@50 | 0.2818 [0.2722, 0.2913] | 0.2978 [0.2888, 0.3072] | +0.0160 [0.0147, 0.0173] |
| Recall@100 | 0.3933 [0.3823, 0.4040] | 0.4215 [0.4116, 0.4319] | +0.0282 [0.0262, 0.0303] |
| NDCG@10 | 0.3890 [0.3786, 0.3994] | 0.4192 [0.4094, 0.4291] | +0.0302 [0.0254, 0.0351] |
| NDCG@50 | 0.3484 [0.3403, 0.3562] | 0.3828 [0.3749, 0.3904] | +0.0344 [0.0316, 0.0372] |
| NDCG@100 | 0.3712 [0.3627, 0.3794] | 0.4075 [0.3999, 0.4151] | +0.0363 [0.0337, 0.0389] |
| AUPRC | 0.2071 [0.2004, 0.2137] | 0.2392 [0.2329, 0.2456] | +0.0321 [0.0299, 0.0343] |

#### Supplementary Results R3. Sample-Classification Results

Tables S4A and S5 report the 250 disjoint-fold and 389 stratified sample-level tasks separately. Across the 250 primary tasks, the Friedman tests rejected equal method ranks for balanced accuracy ($\chi^2_F = 312.80$), AUROC ($\chi^2_F = 373.22$), and AUPR ($\chi^2_F = 377.70$); each test had five degrees of freedom and remained significant after Holm adjustment ($p_{\text{Holm}} < 2 \times 10^{-65}$). The reported Nemenyi comparisons therefore follow significant metric-specific omnibus tests. When task ranks were first averaged within the 137 study accessions, the three Friedman tests remained significant after Holm adjustment ($p_{\text{Holm}} < 2 \times 10^{-47}$); the accession-level Nemenyi critical difference was 0.64. Waypoint uses the common downstream protocol for all 639 tasks. MGM uses the same protocol for all 250 disjoint-fold tasks and for 335 of the 389 sample-level tasks; the other 54 sample-level tasks retain their earlier MGM results because complete genus-profile coverage was unavailable. Pope-MAP uses the common protocol for all 639 tasks.

Figure S4 compares task-weighted and accession-weighted mean scores. Changes were small, with the AUROC and AUPR method ordering preserved.

Table S4: Classification performance for the 250 classification tasks evaluated with disjoint folds. Panel A: Entries are mean ± standard deviation across the task-level cross-validation mean scores. Panel B: accession-equal means for the three newly pretrained conditions, with task-equal means in parentheses. Panel C: paired changes in percentage points, with pointwise 95% accession-bootstrap intervals and Holm-adjusted $p$-values across nine tests. The matched protocol is in Supplementary Methods M1F.

**A. Six-method benchmark**

| Method | Balanced accuracy | AUROC | AUPR |
|---|---|---|---|
| Logistic Regression | $0.633 \pm 0.175$ | $0.707 \pm 0.179$ | $0.653 \pm 0.217$ |
| Random Forest | $0.674 \pm 0.188$ | $0.749 \pm 0.191$ | $0.705 \pm 0.231$ |
| OTU–Taxa | $0.674 \pm 0.190$ | $0.744 \pm 0.189$ | $0.695 \pm 0.229$ |
| Pope-MAP | $0.586 \pm 0.173$ | $0.660 \pm 0.172$ | $0.602 \pm 0.222$ |
| MGM | $0.660 \pm 0.187$ | $0.730 \pm 0.192$ | $0.683 \pm 0.228$ |
| Waypoint | $0.658 \pm 0.186$ | $0.725 \pm 0.191$ | $0.677 \pm 0.230$ |

**B. Matched-pretraining mean scores**

| Pretraining condition | Balanced accuracy | AUROC | AUPR |
|---|---|---|---|
| Paired inputs, joint loss | 0.6771 (0.6752) | 0.7562 (0.7450) | 0.7092 (0.6950) |
| Paired inputs, OTU loss | 0.6762 (0.6726) | 0.7551 (0.7430) | 0.7096 (0.6944) |
| OTU-only inputs, OTU loss | 0.6754 (0.6734) | 0.7559 (0.7439) | 0.7085 (0.6940) |

**C. Paired component effects**

| Added component | Metric | Change (pp) | 95% CI (pp) | Holm $p$ |
|---|---|---|---|---|
| Taxonomy loss | Balanced accuracy | +0.081 | [-0.338, +0.491] | 1.000 |
| | AUROC | +0.104 | [-0.199, +0.405] | 1.000 |
| | AUPR | -0.044 | [-0.372, +0.285] | 1.000 |
| Taxonomy inputs + loss | Balanced accuracy | +0.169 | [-0.249, +0.592] | 1.000 |
| | AUROC | +0.023 | [-0.344, +0.390] | 1.000 |
| | AUPR | +0.069 | [-0.315, +0.469] | 1.000 |
| Taxonomy inputs | Balanced accuracy | +0.087 | [-0.249, +0.438] | 1.000 |
| | AUROC | -0.081 | [-0.408, +0.262] | 1.000 |
| | AUPR | +0.113 | [-0.213, +0.468] | 1.000 |

Table S5: Across-task performance for the 389 classification tasks evaluated with stratified sample-level folds. Entries are mean ± standard deviation across the task-level cross-validation mean scores.

| Method | Balanced accuracy | AUROC | AUPR |
|---|---|---|---|
| Logistic Regression | $0.727 \pm 0.183$ | $0.821 \pm 0.161$ | $0.770 \pm 0.204$ |
| Random Forest | $0.810 \pm 0.179$ | $0.886 \pm 0.145$ | $0.854 \pm 0.184$ |
| OTU–Taxa | $0.801 \pm 0.188$ | $0.871 \pm 0.154$ | $0.835 \pm 0.195$ |
| Pope-MAP | $0.658 \pm 0.214$ | $0.750 \pm 0.182$ | $0.696 \pm 0.232$ |
| MGM | $0.781 \pm 0.190$ | $0.857 \pm 0.160$ | $0.820 \pm 0.200$ |
| Waypoint | $0.774 \pm 0.195$ | $0.848 \pm 0.167$ | $0.811 \pm 0.206$ |

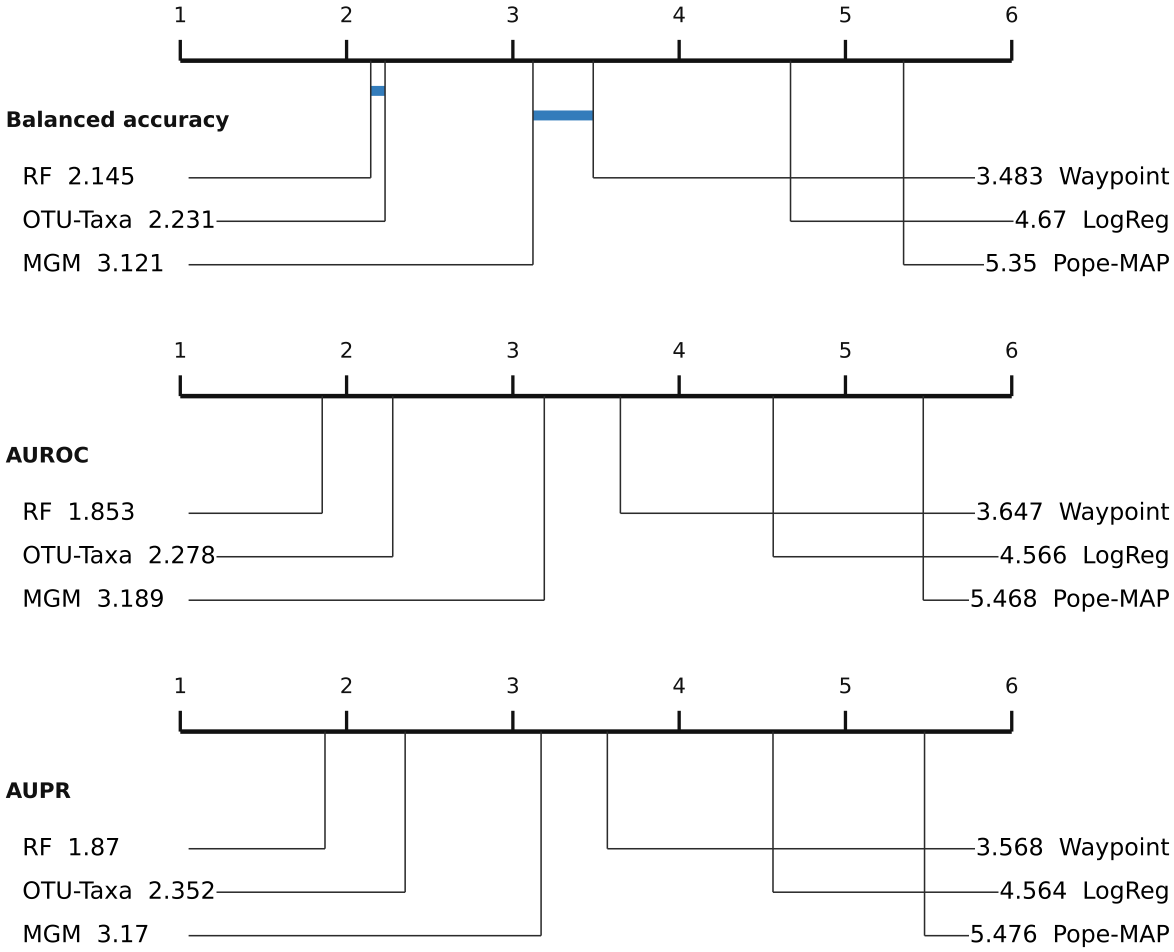


Figure S3: Metric-specific critical-difference summaries for the 389 tasks evaluated with stratified sample-level folds. Lower average rank indicates better performance; the Nemenyi critical difference is 0.38 in each panel.

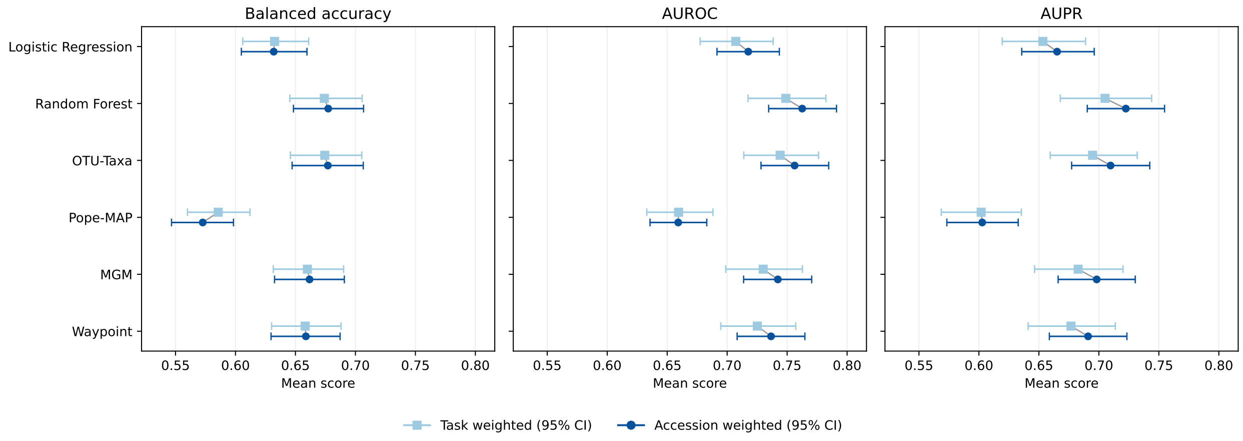


Figure S4: Sensitivity of classification performance to task and accession weighting. Light squares represent equally weighted means across 250 classification tasks. Dark circles represent means calculated first within each study accession and then equally across the 137 accessions. Both summaries use the same fitted models and cross-validation scores. Horizontal bars indicate 95% percentile confidence intervals from 10,000 whole-accession bootstrap resamples, retaining all tasks from each sampled accession together. The same resamples were used for both weighting schemes and all methods and metrics. Changes in mean scores were small; AUROC and AUPR method ordering was preserved, while the closely matched balanced-accuracy means of Random Forest and OTU–Taxa exchanged order.

Table S6: Subject-disjoint performance on the two reference MAP sample-classification tasks. Entries are mean ± standard deviation across the five fixed outer folds. AGP IBD contains $n = 18{,}585$ samples and HMP body site contains $n = 7{,}823$ samples.

| Task | Method | Balanced accuracy | AUROC | AUPR |
|---|---|---|---|---|
| AGP IBD | Logistic Regression | $0.506 \pm 0.130$ | $0.557 \pm 0.158$ | $0.055 \pm 0.011$ |
| AGP IBD | Random Forest | $0.507 \pm 0.052$ | $0.535 \pm 0.171$ | $0.056 \pm 0.016$ |
| AGP IBD | OTU–Taxa | $0.554 \pm 0.077$ | $0.630 \pm 0.114$ | $0.070 \pm 0.019$ |
| AGP IBD | Pope-MAP | $0.513 \pm 0.089$ | $0.574 \pm 0.108$ | $0.060 \pm 0.021$ |
| AGP IBD | MGM | $0.662 \pm 0.131$ | $0.737 \pm 0.130$ | $0.181 \pm 0.193$ |
| AGP IBD | Waypoint | $0.570 \pm 0.062$ | $0.656 \pm 0.077$ | $0.083 \pm 0.015$ |
| HMP body site | Logistic Regression | $0.932 \pm 0.016$ | $0.983 \pm 0.005$ | $0.931 \pm 0.020$ |
| HMP body site | Random Forest | $0.918 \pm 0.008$ | $0.987 \pm 0.002$ | $0.937 \pm 0.007$ |
| HMP body site | OTU–Taxa | $0.948 \pm 0.011$ | $0.989 \pm 0.003$ | $0.961 \pm 0.011$ |
| HMP body site | Pope-MAP | $0.931 \pm 0.013$ | $0.993 \pm 0.001$ | $0.958 \pm 0.009$ |
| HMP body site | MGM | $0.915 \pm 0.011$ | $0.990 \pm 0.003$ | $0.946 \pm 0.011$ |
| HMP body site | Waypoint | $0.936 \pm 0.009$ | $0.991 \pm 0.001$ | $0.956 \pm 0.008$ |

**Matched pretraining controls.** The three pretraining conditions showed similar classification scores across the 250 tasks (Table S4B). Accession-equal paired mean changes were at most 0.169 percentage points in either direction. All nine pointwise confidence intervals included zero and all nine Holm-adjusted $p$-values were 1.000 (Table S4C; Figure S5). These comparisons did not detect significant classification differences between conditions under the matched protocol in Supplementary Methods M1F.

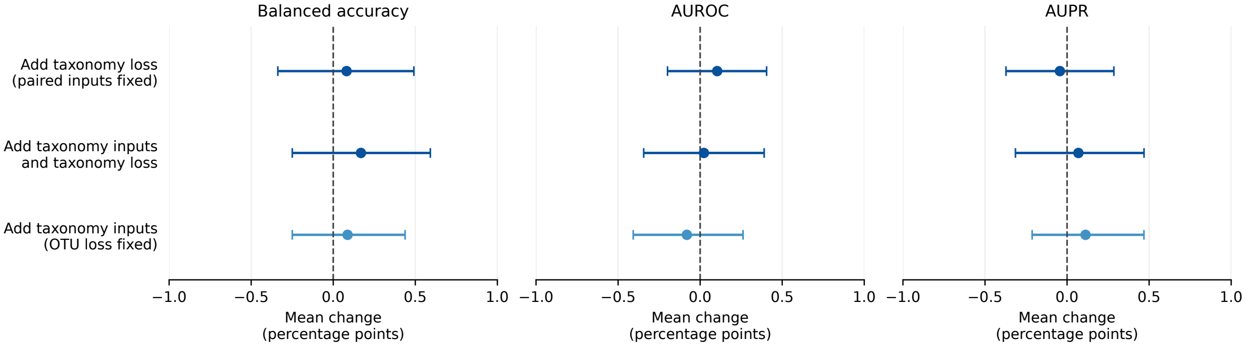


Figure S5: Paired classification effects of taxonomy inputs and supervision. Each point is an accession-equal mean paired change across 137 accessions containing 250 tasks; positive values favor the added component. Bars show pointwise 95% percentile intervals from 10,000 shared whole-accession bootstrap draws, and dashed lines mark zero. Taxonomy loss compares paired inputs with joint loss against paired inputs with OTU loss; taxonomy inputs plus loss compares the joint condition against OTU-only training; taxonomy inputs compares paired inputs with OTU loss against OTU-only training. Table S4C reports the nine Holm-adjusted tests. Supplementary Methods M1F gives the matched protocol.

#### Supplementary Results R4. Sample-Generation Results

Only 19 of 30,000 generated profiles exactly matched a training profile. Across the ten generation replicates, mean (SD) nearest-training-profile Jaccard similarity was 0.2593 (0.0029). Across the 30,000 generated profiles, median nearest-training OTU-set Jaccard similarity was 0.2489, with 90th, 95th, and 99th percentiles of 0.3953, 0.4643, and 0.6250, respectively. Similarity was $\geq 0.50$ for 1,014 profiles (3.38%), $\geq 0.75$ for 92 (0.3067%), $\geq 0.90$ for 23 (0.0767%), and $\geq 0.95$ for 19 (0.0633%). All 19 profiles meeting the last threshold were exact OTU-set matches.

Before no-repeat masking and renormalization, previously emitted OTUs carried mean probability mass 0.1637 (95% profile-bootstrap CI 0.1628–0.1646) across 3,350,132 generation decisions. The decision-level median was 0.1604, and the 95th percentile was 0.3050. Most mass was assigned to a new OTU or the end-of-sample token, although the constraint removed a non-negligible share during decoding.

Table S7: Generation and community-completion benchmarks. Panel A compares the final hierarchy-aware OTU–Taxa checkpoint with global and same-study random training-profile retrieval. Values are means ± sample SD across ten stochastic replicates; OTU–Taxa and global retrieval used 3,000 profiles per replicate, and same-study retrieval covered 2,904 targets (96.8%). Arrows indicate the favorable direction. Panels B and C report community completion on 2,000 held-out test samples as means with 95% percentile-bootstrap confidence intervals from 10,000 sample-level resamples in brackets. Rows labeled "OTU–Taxa minus retrieval" report the paired mean difference and paired bootstrap interval; positive values favor OTU–Taxa.

**A. Unconditional generation and empirical retrieval**

| Metric | OTU–Taxa | Global retrieval | Same-study retrieval |
|---|---|---|---|
| Richness JS ↓ | $0.0165 \pm 0.0016$ | $0.0059 \pm 0.0008$ | $0.0025 \pm 0.0005$ |
| Prevalence Pearson ↑ | $0.9586 \pm 0.0029$ | $0.9689 \pm 0.0019$ | $0.9863 \pm 0.0005$ |
| Co-occurrence Pearson ↑ | $0.9771 \pm 0.0010$ | $0.9854 \pm 0.0009$ | $0.9912 \pm 0.0005$ |
| Co-occurrence RMSE ↓ | $0.0678 \pm 0.0008$ | $0.0345 \pm 0.0009$ | $0.0267 \pm 0.0007$ |

**B. Community-completion recall**

| Method | Recall@10 | Recall@50 | Recall@100 |
|---|---|---|---|
| OTU–Taxa AR completion | 0.101 [0.095, 0.106] | 0.282 [0.272, 0.291] | 0.393 [0.382, 0.404] |
| Sample retrieval | 0.113 [0.108, 0.118] | 0.263 [0.256, 0.270] | 0.322 [0.315, 0.330] |
| OTU–Taxa minus retrieval | -0.012 [-0.018, -0.006] | 0.019 [0.008, 0.030] | 0.071 [0.058, 0.083] |
| Word2vec co-occurrence | 0.064 [0.061, 0.067] | 0.195 [0.189, 0.202] | 0.291 [0.282, 0.299] |
| Learned-embedding co-occurrence | 0.047 [0.044, 0.050] | 0.148 [0.142, 0.153] | 0.223 [0.216, 0.230] |
| Global OTU popularity | 0.015 [0.014, 0.017] | 0.058 [0.054, 0.061] | 0.089 [0.084, 0.094] |

**C. Community-completion ranking**

| Method | NDCG@10 | NDCG@50 | NDCG@100 | AUPRC |
|---|---|---|---|---|
| OTU–Taxa AR completion | 0.389 [0.379, 0.399] | 0.348 [0.340, 0.356] | 0.371 [0.363, 0.379] | 0.207 [0.200, 0.214] |
| Sample retrieval | 0.482 [0.469, 0.494] | 0.412 [0.402, 0.422] | 0.390 [0.381, 0.399] | 0.218 [0.210, 0.225] |
| OTU–Taxa minus retrieval | -0.093 [-0.107, -0.079] | -0.064 [-0.076, -0.052] | -0.018 [-0.030, -0.007] | -0.010 [-0.020, -0.001] |
| Word2vec co-occurrence | 0.293 [0.284, 0.303] | 0.260 [0.253, 0.267] | 0.278 [0.272, 0.285] | 0.140 [0.136, 0.145] |
| Learned-embedding co-occurrence | 0.222 [0.214, 0.230] | 0.199 [0.193, 0.205] | 0.215 [0.209, 0.221] | 0.101 [0.098, 0.105] |
| Global OTU popularity | 0.071 [0.066, 0.076] | 0.067 [0.063, 0.071] | 0.076 [0.072, 0.080] | 0.030 [0.029, 0.032] |

**Sequential prediction comparison.** Mean hidden-member recall was 24.49% for fixed-prefix ranking, 25.29% for sequential greedy prediction, 15.13% for frozen sampling, and 13.44% for sequential sampling (Figure S6). Greedy continuation improved recall over fixed-prefix ranking by 0.80 percentage points (95% paired bootstrap CI, 0.41–1.20), whereas sequential sampling performed worse than frozen sampling. Retrieval achieved higher recall, at 29.66%. These comparisons show that sequential prediction gives a small improvement with greedy selection, while probabilistic sampling does not improve recovery under the tested settings.

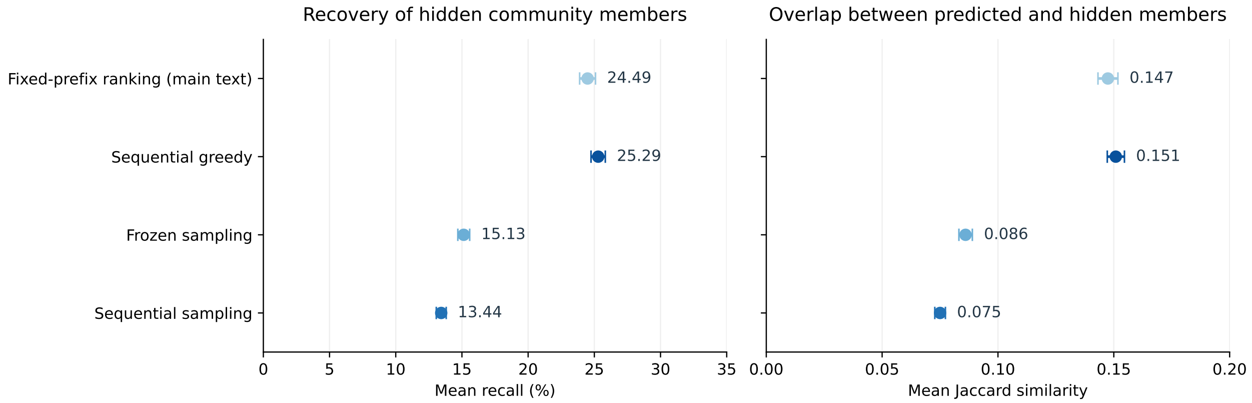


Figure S6: Comparison of four OTU–Taxa prediction procedures after observing the first half of each abundance-ranked profile. The left panel shows the percentage of hidden OTUs recovered; the right shows Jaccard similarity between predicted and hidden OTU sets. Points are means across 2,000 profiles, with 95% confidence intervals from 10,000 profile-bootstrap resamples. Sampling results average three seeds within each profile. Every procedure outputs exactly the number of hidden OTUs, excluding observed and previously selected OTUs. The "main text" label identifies the fixed-prefix prediction procedure, evaluated here with this shared output-length budget rather than the fixed ranking cutoffs used in the main benchmark.

#### Supplementary Results R5. Disease-Ecology Results

Supplementary Figure S7 shows two contrasting task-level abundance-rank readouts: strong positive model–count agreement in a plant drought comparison and systematic inversion in a lake-versus-stream comparison.

The aggregate analysis summarizes task-level model–count agreement after restricting each contrast to nested subsets of count-selected genera defined by pooled observed relative abundance. Across the 78 retained contrasts from 74 study accessions, the conditional mean AUROC was 0.5884 (95% study-cluster bootstrap interval 0.5359–0.6417). The corresponding permutation null was centred at 0.4998, and none of the 10,000 permutation replicates reached the observed mean (one-sided $p = 0.00010$). Mean AUROC reached 0.6727 for the top half and 0.7179 for the top quarter. Relative to all selected genera, the corresponding increases were 0.0843 (95% study-cluster interval 0.0679–0.1005) and 0.1294 (0.1058–0.1544).

Supplementary Figure S8 complements this task-level analysis by examining whether genera that appear earlier in observed abundance-ranked profiles contribute more frequently to concordant model–count AUROC orderings. The association was positive but modest (Spearman $\rho = 0.1743$; Pearson $r = 0.1851$). BugSigDB concordance in Supplementary Table S8 provides an additional exploratory comparison with literature-derived disease directions.

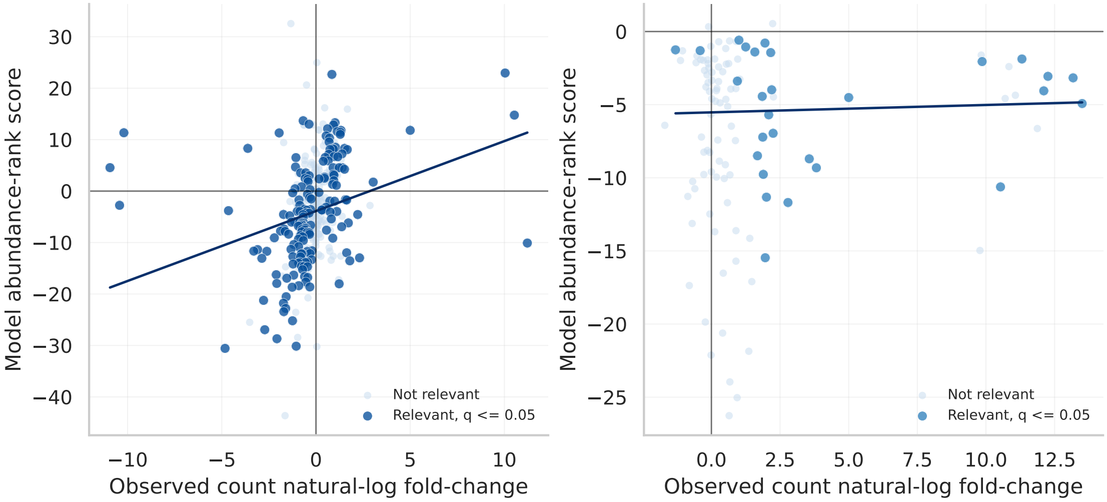


Figure S7: Model–count agreement in two generative abundance-rank readouts. The left panel shows the plant drought comparison and the right panel shows lake versus stream water. Each point is a genus; the horizontal axis is its observed count logFC and the vertical axis is its model abundance-rank score. Dark points pass the task-specific count-based threshold $q \leq 0.05$, light points are other shared genera, and lines are least-squares fits through the selected genera.

Table S8: Agreement between the model abundance-rank readout, observed count logFC, and pooled BugSigDB IBD-related directions on the common BugSigDB genera shared by the model and count tables for each cohort. AUROC values compare each readout with the sign of the BugSigDB net direction.

| Cohort | $n$ genera | Model abundance-rank AUROC | Observed count-logFC AUROC |
|---|---|---|---|
| AGP | 122 | 0.674 | 0.406 |
| Halfvarson | 72 | 0.770 | 0.808 |

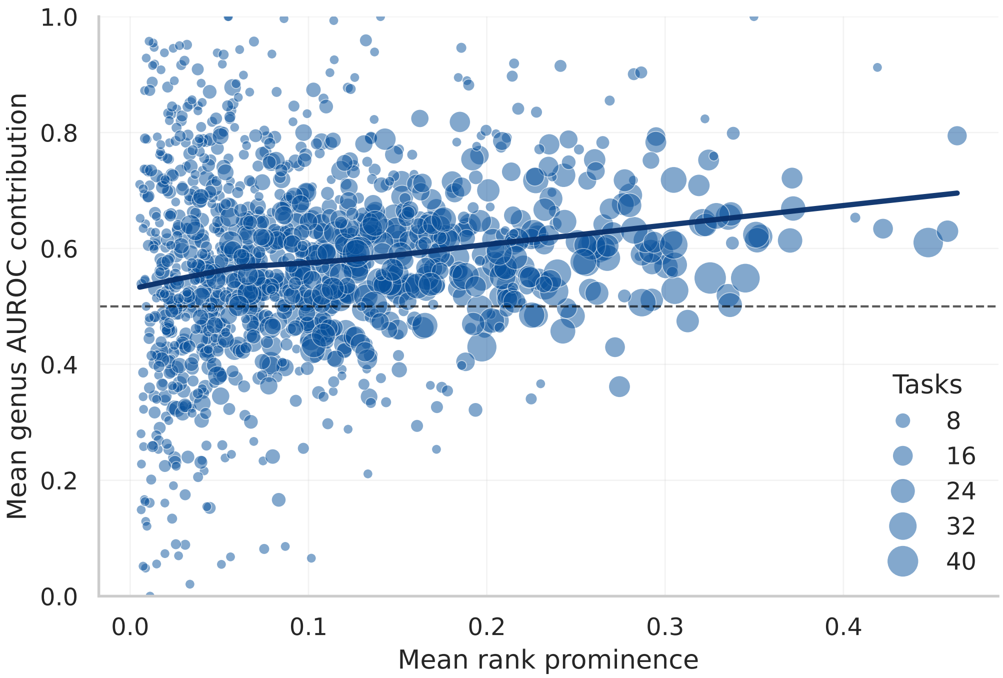


Figure S8: Relationship between observed abundance-rank prominence and genus contribution to the model–count AUROC readout. Each point is one genus summarized across the disease/ecology tasks in which it was selected by the task-specific count-based FDR filter. Rank prominence is defined as 1− normalized observed rank fraction, so larger values indicate genera that appear earlier in the abundance-ranked sample profile. Mean genus AUROC contribution is the average fraction of AUROC pairwise comparisons involving that genus in which the model–count ordering agrees with the observed direction. The positive but modest association (Spearman $\rho = 0.1743$, Pearson $r = 0.1851$) indicates that genera appearing earlier in observed abundance-ranked profiles contribute slightly more often to concordant model–count pairwise orderings.

#### Supplementary Results R6. Context-Based Taxonomy-Assignment Recovery

Figures S9 and S12 show the first rank at which SINTAX- and QIIME2-aligned context predictions diverge from their reference paths. Across OTUs, mean rankACC was 0.4561 in the SINTAX label space and 0.4421 in the QIIME2 label space. These values are the OTU-count-weighted means across the deepest-reference-rank strata shown in the error-origin matrices. Per-rank agreement decreased toward finer taxonomic levels, with complete-path exact matches for 10.1% and 10.7% of the SINTAX- and QIIME2-aligned OTUs, respectively (Table S9).

Table S9: Per-rank and complete-path agreement of OTU-level majority predictions with intact-sequence classifier assignments in the SINTAX and QIIME2 label spaces. Agreement is shown with a 95% Wilson confidence interval across evaluable OTUs. The number of evaluable OTUs varies at finer ranks because species labels and some intermediate annotations are not available for every reference path.

| | SINTAX label space | | QIIME2 label space | |
|---|---|---|---|---|
| Rank | $n$ | Agreement [95% CI] | $n$ | Agreement [95% CI] |
| Kingdom | 5,666 | 0.991 [0.988, 0.993] | 4,168 | 0.992 [0.988, 0.994] |
| Phylum | 5,666 | 0.602 [0.589, 0.614] | 4,168 | 0.589 [0.574, 0.603] |
| Class | 5,660 | 0.450 [0.437, 0.463] | 4,161 | 0.436 [0.421, 0.451] |
| Order | 5,620 | 0.406 [0.393, 0.419] | 4,153 | 0.388 [0.373, 0.403] |
| Family | 5,538 | 0.301 [0.289, 0.313] | 4,082 | 0.309 [0.295, 0.323] |
| Genus | 5,666 | 0.158 [0.148, 0.167] | 4,168 | 0.188 [0.176, 0.200] |
| Species | 557 | 0.009 [0.004, 0.021] | 872 | 0.018 [0.011, 0.030] |
| Complete path | 5,666 | 0.101 [0.094, 0.109] | 4,168 | 0.107 [0.098, 0.117] |

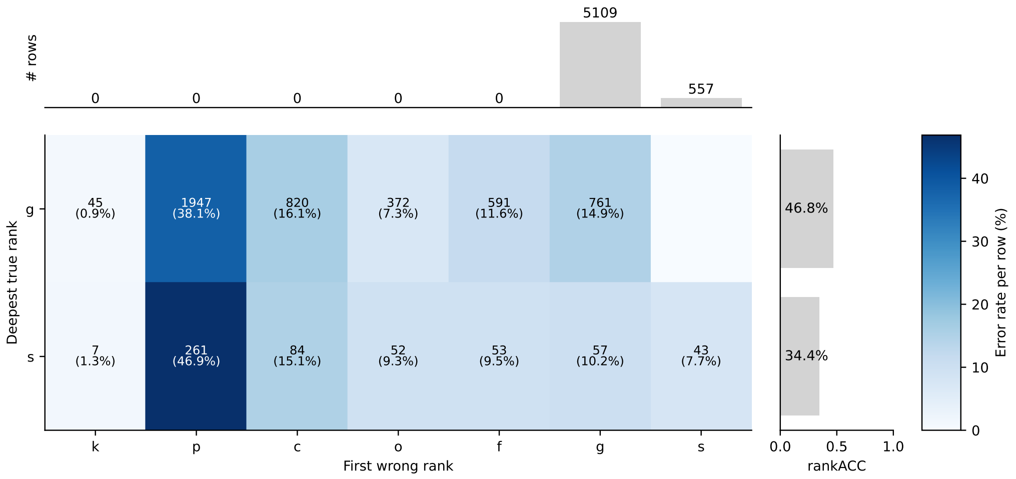


Figure S9: Error-origin matrix for SINTAX-aligned context-based taxonomy-assignment recovery. Rows indicate the deepest evaluated reference rank, and columns indicate the first rank at which the context-only OTU–Taxa prediction diverges from the reference path. This view helps separate shallow mistakes, where the context prediction diverges near the root, from deeper mistakes in which the prediction remains correct through several taxonomic ranks.

Figures S10 and S11 characterize how context-based taxonomy-assignment recovery depends on the biological sample and on the two input channels. The global-prior and shuffled-sample conditions assess sample specificity, whereas the OTU-only and taxonomy-only conditions quantify how much signal is retained through either channel of the same pretrained representation. The visible-context analysis varies the amount of observed prefix context.

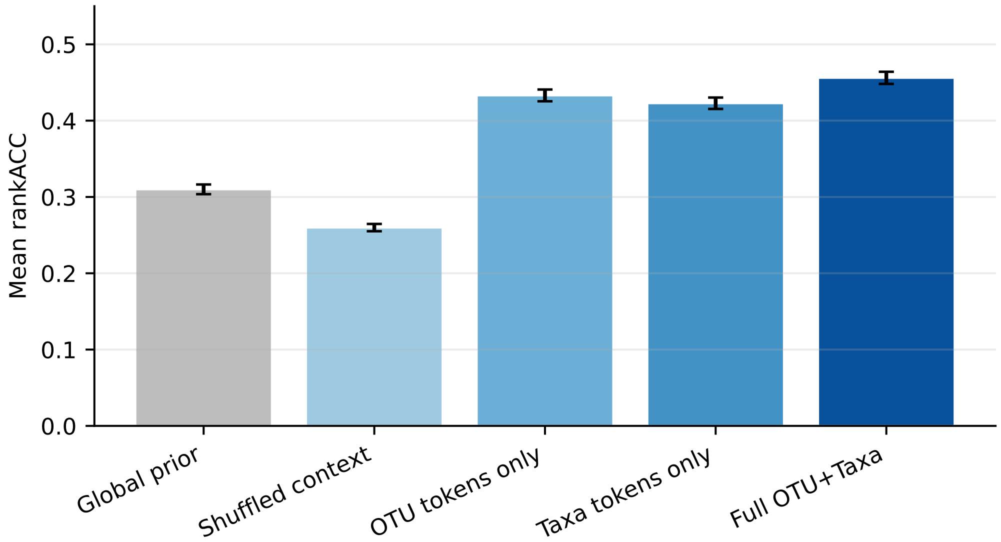


Figure S10: SINTAX-aligned context-ablation results for context-based taxonomy-assignment recovery. Bars report mean OTU-level rankACC after rank-specific majority aggregation across occurrences: 0.310 for the global prior, 0.260 for shuffled sample context, 0.433 for OTU-only context, 0.423 for taxonomy-only context, and 0.456 for full OTU–taxonomy context.

The matched QIIME2-labeled model reached mean OTU-level rank accuracy relative to QIIME2 assignments of 0.442 (95% CI, 0.433–0.451) across 4,168 OTUs.

[5] Tackmann J, Arora N, Schmidt TSB, Rodrigues JFM, von Mering C. Ecologically informed microbial biomarkers and accurate classification of mixed and unmixed samples in an extensive

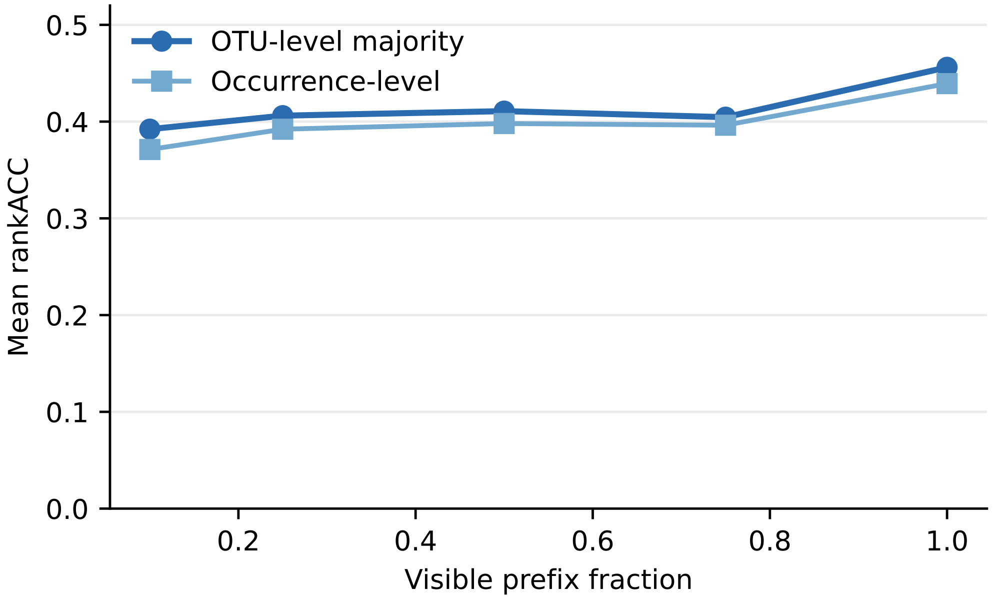


Figure S11: SINTAX-aligned visible-context fraction analysis for context-based taxonomy-assignment recovery. Mean OTU-level rankACC was 0.392, 0.406, 0.411, 0.405, and 0.456 when the first 10%, 25%, 50%, 75%, and 100% of the available real prefix positions were retained, respectively.

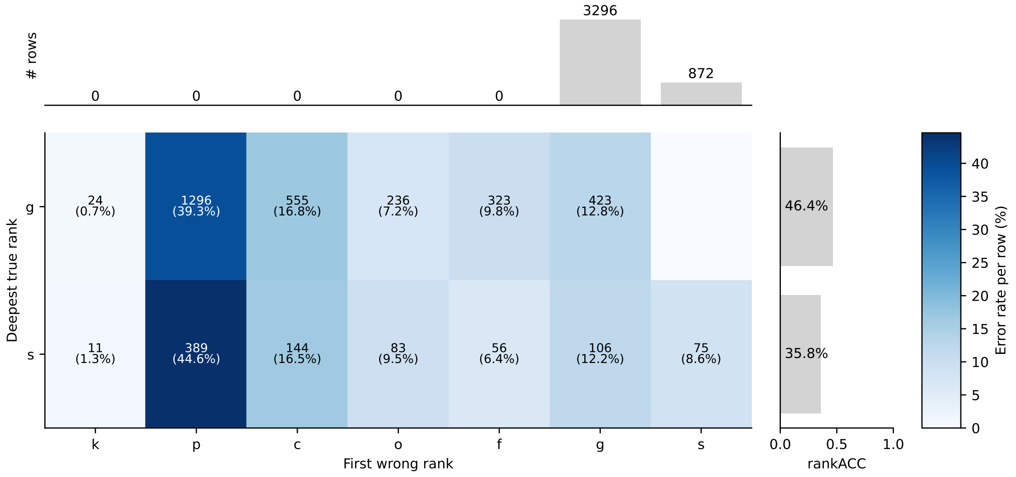


Figure S12: Error-origin matrix for QIIME2-aligned context-based taxonomy-assignment recovery. Rows indicate the deepest evaluated reference rank, and columns indicate the first rank at which the context-only OTU–Taxa prediction diverges from the reference path.

cross-study of human body sites. Microbiome. 2018;6(1):192. https://doi.org/10.1186/s40168-018-0565-6.
